# Cortical Face-Selective Responses Emerge Early in Human Infancy

**DOI:** 10.1101/2021.12.04.471085

**Authors:** Heather L. Kosakowski, Michael A. Cohen, Lyneé Herrera, Isabel Nichoson, Nancy Kanwisher, Rebecca Saxe

## Abstract

In human adults, multiple cortical regions respond robustly to faces, including the occipital face area (OFA) and fusiform face area (FFA), implicated in face perception, and the superior temporal sulcus (STS) and medial prefrontal cortex (MPFC), implicated in higher level social functions. When in development does face selectivity arise in each of these regions? Here, we combined two awake infant functional magnetic resonance neuroimaging (fMRI) datasets to create a sample size twice the size of previous reports (n=65 infants, 2.6-9.6 months). Infants watched movies of faces, bodies, objects, and scenes while fMRI data were collected. Despite variable amounts of data from each infant, individual subject whole-brain activations maps revealed a significant response to faces compared to non-face visual categories in the approximate location of OFA, FFA, STS, and MPFC. To determine the strength and nature of face selectivity in these regions, we used cross-validated functional region of interest (fROI) analyses. Across this larger sample size, face responses in OFA, FFA, STS, and MPFC were significantly greater than responses to bodies, objects, and scenes. Even the youngest infants (2-5 months) showed significantly face-selective responses in FFA, STS, and MPFC, but not OFA. These results demonstrate that face selectivity is present in multiple cortical regions within months of birth, providing powerful constraints on theories of cortical development.

**Significance Statement:** Social cognition often begins with face perception. In adults, several cortical regions respond robustly to faces, yet little is known about when and how these regions first arise in development. To test whether face selectivity changes in the first year of life, we combined two datasets, doubling the sample size relative to previous reports. In the approximate location of the fusiform face area (FFA), superior temporal sulcus (STS), and medial prefrontal cortex (MPFC) but not occipital face area (OFA), face selectivity was present in the youngest group. These findings demonstrate that face-selective responses are present across multiple lobes of the brain very early in life.

## INTRODUCTION

Faces are highly salient visual and social features of our environment. In human adults, many cortical regions show robust and selective responses to faces (Dai and Scherf, 2023; Haxby et al., 2000). We focus on four such regions: the occipital face area (OFA) in the inferior occipital gyrus (Gauthier et al., 2000), the fusiform face area (FFA) in the fusiform gyrus (Kanwisher et al., 1997), and regions in the superior temporal sulcus (STS) and the medial prefrontal cortex (MPFC). Here we ask when in development each of these regions first respond selectively to faces.

OFA and FFA are both visual regions, responding robustly to visually presented faces and much less to any other visual category. OFA is anatomically posterior to FFA and appears to encode face features or parts (Henriksson et al., 2015). FFA, on the other hand, appears to encode the presence and identity of a face more holistically (Grill-Spector et al., 2004; Yovel and Kanwisher, 2005). Individual neurons within FFA are highly face selective (Axelrod et al., 2019; Khuvis et al., 2021) and electrically stimulating this area can distort or create face percepts (Jonas et al., 2018; Parvizi et al., 2012; Rangarajan et al., 2014; Schalk et al., 2017).

By contrast, STS and MPFC contain regions that respond robustly to faces, but these responses are modulated by social context, and the same regions also respond to socially relevant stimuli that are not faces. A region in STS is face-selective when compared to other visual categories, and preferentially responds to socially relevant movements of faces, like facial expressions and shifts of eye gaze (Pitcher et al., 2011), but also responds to human voices (Deen et al., 2020). Similarly, a region in MPFC is face-selective on visual tasks, but the response to faces is influenced by the faces’ social attributes such as moral goodness and attractiveness (Cheng et al., 2022; LaBar, 2003; O’Doherty et al., 2003). The same region in MPFC also responds to socially-relevant verbal narratives (Kosakowski et al., 2022b). In sum, in adults OFA, FFA, STS, and MPFC all have face-selective responses, though plausibly perform different visual and social functions.

Initial fMRI studies revealed substantial changes in the extent and magnitude of face-selective responses throughout childhood and early adolescence, suggesting that face-selectivity is slow to develop (Cohen Kadosh et al., 2013, 2011; Feng et al., 2022; Golarai et al., 2015, 2009, 2007; Haist et al., 2013; Joseph et al., 2011; Natu et al., 2016; Nordt et al., 2021; Peelen et al., 2009; Scherf et al., 2007; Tian et al., 2021). On the other hand, human infants have at least modest preferential responses to faces in the approximate location of OFA, FFA, STS, and MPFC (Deen et al., 2017; Ichikawa et al., 2010; Lisboa et al., 2020b; Lloyd-Fox et al., 2017, 2011a, 2009a; Powell et al., 2018a; Tzourio-Mazoyer et al., 2002). Most of these prior studies did not measure responses to other visual categories, to establish whether the responses were face selective. One recent study reported face-selective responses in FFA in infants, but did not investigate STS or MPFC (Kosakowski et al., 2022a). Additionally, because of small sample sizes, prior studies could not resolve when face-selective responses in these regions first appears, within the first year of life. Thus, it remains an open question when each of these cortical regions first shows face-selective responses (Scott and Arcaro, 2023).

Functional MRI (fMRI) is the only neuroimaging method that has the coverage and spatial resolution to measure neural responses simultaneously in OFA, FFA, STS, and MPFC. A substantial challenge for fMRI with infant populations is that fMRI requires the participant to be still for long periods of time during data acquisition making awake infant fMRI studies rare (Ellis et al., 2020).

For the current study, we combined two fMRI datasets that were collected (Figure 1a) while infants watched dynamic videos of faces (Figure 1b), bodies, objects, and scenes (Figure 1c). In this combined dataset, infants ranged in age from 2 to 9 months (n=65; Figure 1d). As a result, the oldest infants had three times as much post-birth experience as the youngest ones. Thus, these data allow us to test whether, and when, cortical face selectivity emerges in OFA, FFA, STS, and MPFC in the first year of human infants’ life.

**Figure 1.**
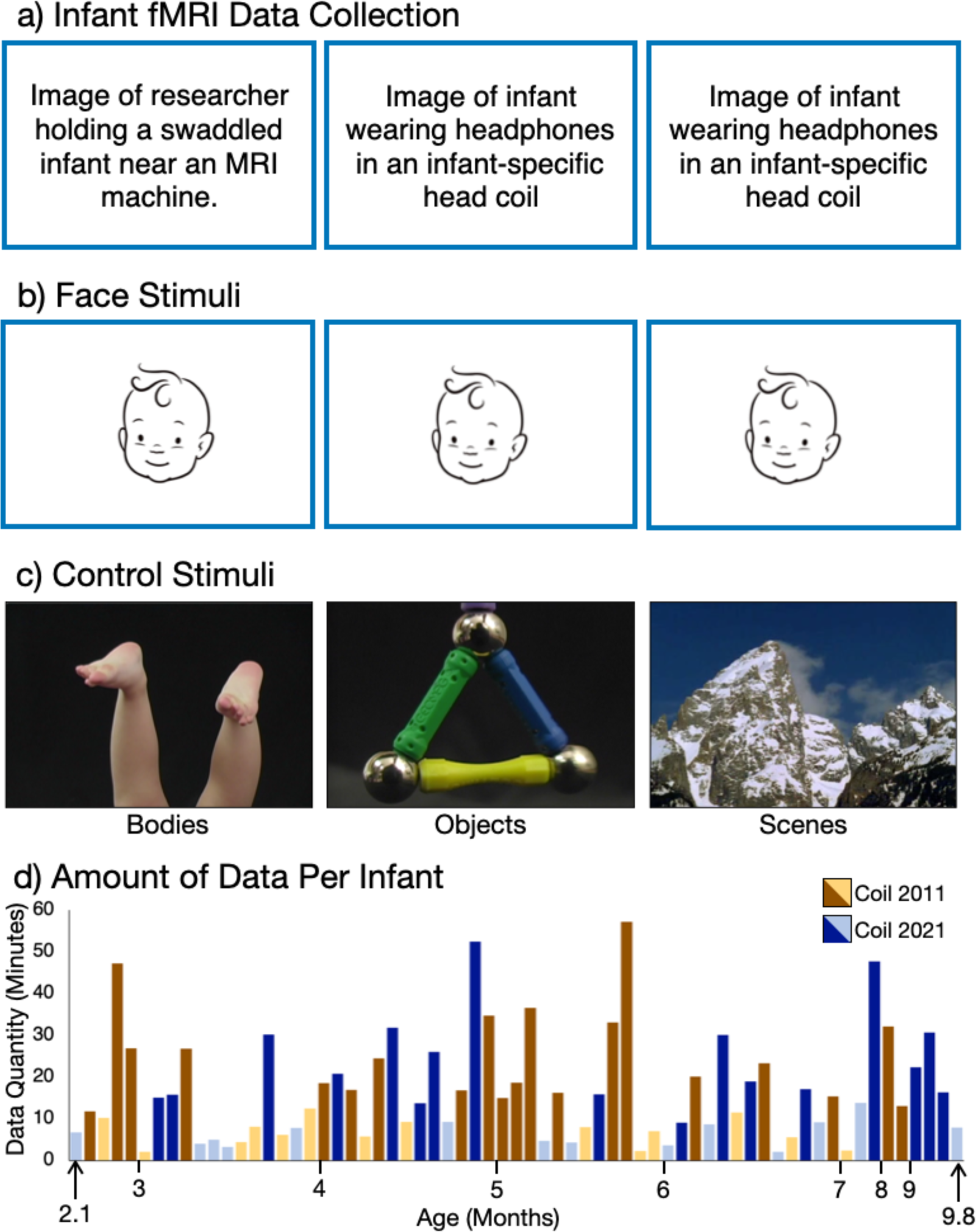
Awake Infant fMRI Scanning and Stimuli. (a) For each MRI visit we swaddled the infant, applied hearing protection, and placed the infant in a custom 32-channel infant head coil. Movies were projected in a mirror over infants’ eyes. Photo credit for images in the middle and right to Caitlin Cunningham Photography. Photo credit for the image on the left to Kris Brewer. (b) Example frames from dynamic face stimuli. (c) Example frames from dynamic body, object, and scene stimuli. (d) Infants that had enough usable data to compute a whole-brain contrast image are plotted on the x-axis, ordered by age in months. Data collected using Coil 2011 is indicated in brown, data collected using Coil 2021 is indicated in blue. Darker color indicates inclusion in fROI analyses (see Methods for inclusion criteria).

## METHODS

### Infant fMRI Data

To increase our ability to measure age differences, we combined analyses of data that were collected using two different infant head coils. One group of infants (n=31) was scanned from June 2016 to July 2019 with one coil (Coil 2011 described below; (Keil et al., 2011)) with a sinusoidal acquisition sequence (Zapp et al., 2012) on a 3T Siemens Trio Scanner. A second group of infants (n=56) was scanned from July 2019 to February 2020 with a different coil (Coil 2021 described below (Ghotra et al., 2021)) using a higher resolution acquisition sequence (see Data Collection) on a 3T Siemens Prisma Scanner. Each infant was scheduled for up to 8 visits. Any usable data that were collected within a 30-day window on the same coil with the same acquisition sequence were analyzed as a single session (see Data Selection). In total, we had 74 sessions from 61 individuals (2.0-9.8 months, mean=5.2 months). For each session, we collected 4.30-112.70 minutes of data (mean=34.92 min, s.d.=22.45 min). To be eligible for analysis, we identified segments of usable data with less than 2mm/radians of frame-to-frame displacement, resulting in 65 usable sessions (2.1-9.8 months, mean=5.3) from 53 unique individuals with 2.05-57.25 minutes of usable data per session (Figure 1d; mean=16.99, s.d.=12.82). Of these, 49 sessions from 46 unique individuals had enough data to be included in whole brain random effects analyses and 37 sessions from 33 unique individuals met the inclusion criteria for functional region of interest (fROI) analyses (Figure 1d).

### Participants

Infants (n=86; 2.0-11.9 months; mean=5.4 months; 41 female) were recruited at a location that will be identified if the article is published through word-of-mouth, fliers, and social media. These data have been previously reported in (Kosakowski et al., 2022a), mainly reporting data from Coil 2021 and measuring responses only in visually responsive category-selective regions. Here, the data were reanalyzed with a focus on face-responsive regions across the cerebral cortex and combining data from both Coil 2021 and Coil 2011 (Keil et al., 2011) to create a larger sample. Usable data (see data selection) were collected from 65 infants (2.6-11.9 months; 24 female; 31 from Coil 2021 and 34 from Coil 2011). Parents of participants were provided parking or reimbursed travel expenses. Participants received a small compensation for each visit and, when possible, printed images of their brain. Parents of participants provided informed consent and all protocols were approved by the Institutional Review Board at author university.

### Experimental Paradigms

*Paradigm 1*. Infants watched videos of faces (Figure 1b), bodies, objects, and scenes (Figure 1c) (Pitcher et al., 2011). A colorful, curvy, abstract baseline was used to maintain infants’ attention during baseline blocks. Videos were selected to be categorically homogeneous within blocks and heterogeneous between blocks. Each block was 18s and was composed of six 3s videos from the same category. Face videos showed one child’s face on a black background. Object videos showed toys moving. Body videos showed children’s hands or feet on a black background. Scene videos showed natural environments. Baseline blocks were also 18s and consisted of six 3s videos that featured abstract color scenes such as liquid bubbles or tie-dyed patterns. Block order was pseudo-random such that all blocks played once prior to playing again. Videos played continuously for as long as the infant was content, paying attention, and awake.

*Paradigm 2.* Infants watched videos from the same five conditions as in *Paradigm 1*. However, the videos were shortened to 2.7s and interleaved with still images from the same category (but not drawn from the videos) presented for 300ms. All blocks were 18s and included 6 videos and 6 images. Video and image order were randomized within blocks and block order was pseudorandom by category. *Paradigm 2* contained one additional block depicting hand-object interactions which was not included in the present analysis.

### Data Collection

Infants were swaddled if possible (Figure 1a). A parent or researcher went into the scanner with the infant while a second adult stood outside the bore of the scanner. Infants heard lullabies (https://store.jammyjams.net/products/pop-goes-lullaby-10) for the duration of the scan. For data collected with *Coil 2011*, lullabies were played over a loudspeaker into the scanning room. For data collected with *Coil 2021*, lullabies were played through custom infant headphones (Figure 1a).

*Coil 2011.* For data collected with *Coil 2011* we used a custom 32-channel infant coil designed for a Siemens Trio 3T scanner (Keil et al., 2011) and a quiet EPI with sinusoidal trajectory (Zapp et al., 2012) with 22 near-axial slices (repetition time, TR=3s, echo time, TE=43 ms, flip angle=90°, field of view, FOV=192 mm, matrix=64×64, slice thickness=3 mm, slice gap=0.6 mm). The sinusoidal acquisition sequence caused substantial distortions in the functional images.

*Coil 2021.* Infants wore custom infant MR-safe headphones. Infant headphones attenuated scanner noises and allowed infants to hear the lullabies. An adjustable coil design (Ghotra et al., 2021) increased infant comfort and accommodated headphones as well as a variety of head sizes (Figure 1a). The new infant coil and infant headphones designed for a Siemens Prisma 3T scanner enabled the use of an EPI with standard trajectory with 44 near-axial slices (repetition time, TR=3s, echo time, TE=30ms, flip angle=90°, field of view, FOV=160 mm, matrix=80×80, slice thickness=2 mm, slice gap=0 mm). Six infants had data collected using a different EPI with standard trajectory with 52 near-axial slices (repetition time, TR=2s, echo time, TE=30ms, flip angle=90°, field of view, FOV=208 mm, matrix=104×104, slice thickness=2 mm, slice gap=0 mm). Functional data collected with *Coil 2021* were less distorted than data collected with *Coil 2011*.

### Data Selection (Subrun Creation)

To be included in the analysis, data had to meet criteria for low head motion (Deen et al., 2017; Kosakowski et al., 2022a). Data were cleaved between consecutive timepoints with more than 2 radians or mm of frame-to-frame displacement, creating *subruns*, which had to contain at least 24 consecutive low-motion volumes to be included in further analysis. All volumes included in a subrun were extracted from the original run data and combined to create a new Nifti file for each subrun.

Paradigm files indicating which condition occurred at each time point were similarly updated for each subrun. Volumes with greater than 0.5 radians or mm of frame-to-frame displacement from the previous or following volume were scrubbed (i.e., removed) from all analyses. Data collected within a 30-day window from a single subject were analyzed as one session. Figure 2a shows the amount of data collected and the amount of data included in the subruns, prior to scrubbing, for each participant session. The total amount of data initially collected was negatively correlated with age (*r*=-0.26, p=0.003), and the proportion of data we retained per session was positively correlated with age (Figure 2b; *r*= 0.39, *p*<0.001). That is, older infants had shorter sessions, and tended to be still for greater proportions of those sessions. Younger infants were more likely to move and required longer sessions to produce the same eventual yield of usable data.

**Figure 2.**
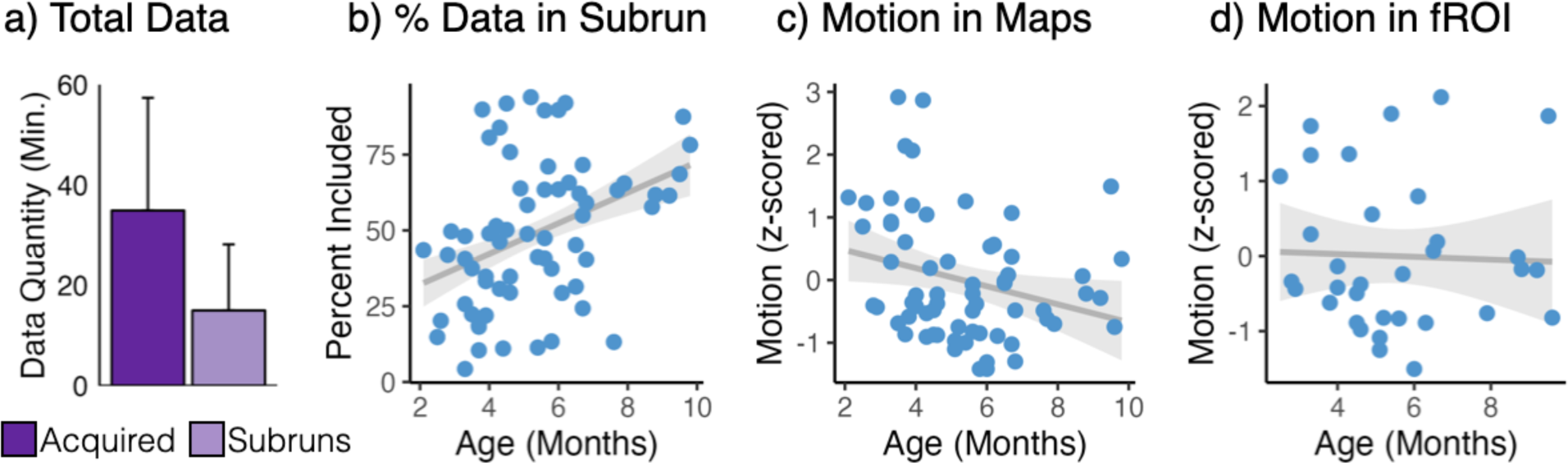
Data Quality Metrics for Awake Infant fMRI. (a) Variable amounts of data were acquired from each 30-day participant session (dark purple) and included in subruns (light purple). Bars indicate mean minutes of data per participant session, error bars are standard deviation. (b) The amount of data included in subruns for a participant session was correlated with age. (c) For participants that had enough data to compute whole-brain contrast maps, the proportion of motion (number of volumes with greater than 0.5 mm or radians of frame-to-frame displacement) was positively correlated with age. (d) For participants with enough data to be included in fROI analyses, motion was not correlated with age.

Participants had to have at least 5 minutes of low-motion data to be included in whole brain analyses. For each participant, only one session was included in each RFX analysis (which were run separately for Coil 2011 and Coil 2021). For fROI analyses, subruns were combined or split, as necessary, to create subruns with at least 96 volumes each. Subruns were designed to have approximately the same number of volumes, within participant (see below for more information). To be included in the fROI analysis, participants had to have at least two subruns (one to choose voxels and the other to extract independent response magnitudes from the selected voxels).

### fMRI Data Preprocessing

Each subrun was processed individually. First, an individual functional image was extracted from the middle of the subrun to be used for registering the subruns to one another for further analysis. Then, each subrun was motion corrected using FSL MCFLIRT. If more than 3 consecutive images had more than 0.5 mm or 0.5 radians of motion, there had to be at least 7 consecutive low-motion volumes following the last high-motion volume for those volumes to be included in the analysis. Additionally, each subrun had to have at least 24 volumes after accounting for motion and timepoints when the infants appeared to be asleep (e.g., with eyes closed). Functional data were skull-stripped (FSL BET2), intensity normalized, and spatially smoothed with a 3mm FWHM Gaussian kernel (FSL SUSAN).

### Data Registration

All subruns were aligned within subjects and then each subject was registered to a standard template. First, the middle image of each subrun was extracted and used as an example image for registration. If the middle image was corrupted by motion or distortion, a better image was selected as the example image. The example image from the middle subrun of the first visit with usable data was used as the target image. All other subruns from each subject were registered to that subject’s target image using FSL FLIRT. The target image for each subject was then registered to a template image using FSL FLIRT. For data collected with *Coil 2011*, the template image was taken from (Deen et al., 2017). For data collected with *Coil 2021*, the template image was taken from (Kosakowski et al., 2022a). Given the distortion of the images and the lack of an anatomical image for each subject, traditional registration tools do not effectively register infant data between subjects. As such, we attempted to register each image using a rigid, an affine, and a partial affine registration with FSL FLIRT. The best image registration was selected by eye from the three options and manually tuned using the FreeSurfer GUI for the best possible data alignment. Each image took between 2 and 8 hours of human labor to register. Images collected with *Coil 2021* were transformed to the anatomical space of the template image for visualization.

### Subject-Level Beta and Contrast Maps

Functional data were analyzed with a whole-brain voxel-wise general linear model (GLM) using custom MATLAB scripts. The GLM included 4 condition regressors (faces, scenes, bodies, and objects), 6 motion regressors, a linear trend regressor, and 5 PCA noise regressors. PCA noise regressors are analogous to GLMDenoise (Kay et al., 2013). Condition regressors were defined as a boxcar function for the duration of each condition block (18s). Infant inattention or sleep was accounted for using a single nuisance (‘sleep’) regressor. The sleep regressor was defined as a boxcar function with a 1 for each TR the infant was not looking at the stimuli, and the corresponding TR was set to 0 for all condition regressors. Boxcar condition and sleep regressors were convolved with an infant hemodynamic response function (HRF) that is characterized by a longer time to peak and deeper undershoot compared to the standard adult HRF (Arichi et al., 2012). Next, data and all regressors except PCA noise regressors were concatenated across subruns. PCA noise regressors were computed across concatenated data and beta values were computed for each condition in a whole-brain voxel-wise GLM. Subject-level contrast maps to test for face-selective responses were computed as the difference between the face beta and the average of all non-face (i.e., bodies, objects, and scenes) betas for each voxel using in-house MATLAB code.

### Group Random Effects Analysis

Due to variable distortions in the BOLD images across participants, and lack of a T1 and/or T2 image from most infants, data registration across participants is imperfect. Additionally, the sequences used with each coil created very different patterns of spatial distortion, so we conducted separate group random effects analyses for data collected from each coil. First, subject-level contrast difference (faces – non-face) maps were transformed to coil-specific template space. Group RFX analyses were performed using FreeSurfer mri_concat and FreeSurfer mri_glmfit. In the dataset used for whole-brain analyses, the amount of motion (the number of scrubbed volumes divided by the number of total volumes) was negatively correlated with age (Figure 2c; *r*=-0.27, *p*= 0.03).

### Functional Region of Interest (fROI) Analysis

Group RFX analyses are imperfect because they rely on high-quality registrations to a common template across subjects and they do not respect idiosyncratic anatomical and functional differences across individuals. Thus, to determine if cortical responses are face-selective, we utilized an fROI approach. Using fROI analyses enables us to (1) account for individual anatomical variability (Saxe et al., 2006), (2) more rigorously characterize responses using a cross-validation procedure (Nieto-Castañón and Fedorenko, 2012), and (3) requires high-quality within-subject registrations but is more tolerant of imperfect across-subject registrations.

We account for the variable amount of data in each subrun for each subject (n=37 sessions, 33 unique individuals) and the impact this could have on reliable parameter estimates from the GLM by first combining or splitting subruns. This allowed us to approximately equate the amount of data across subruns within each subject. For example, if a subject had three subruns and the first had 30 volumes, the second had 75 volumes and third had 325 volumes, then we concatenated the first two subruns to create one subrun and we split the third subrun into 3 resulting in a total of four subruns with approximately 100 volumes each. For data included in fROI analyses, motion and age were not correlated (Figure 2d; *r*=-0.04, *p*=0.8). Thus, the fROI analyses allow for a measurement of age effects that is not confounded with motion.

To constrain search areas for voxel selection, we used anatomically defined parcels transformed to subject-specific space. Due to the distortions in the *Coil 2011* dataset, we opted to use larger parcels than the FFA and OFA parcels used in (Kosakowski et al., 2022a). We created large parcels that extended well beyond the boundaries of face regions in the inferorior occipiatal gyrus (IOG, the approximate locations of OFA), and ventral temporal cortex (VTC, the approximate location of FFA), STS, MPFC using the Glasser atlas (Glasser et al., 2016). The large OFA parcel included Glasser areas LO1, LO2, LO3, V4, V4t and PIT and the large FFA parcel included Glasser areas VMV1, VMV2, VMV3, VVC, PHA1, PHA2, PHA3, and FFC. For the MPFC parcel we used Glasser areas p24, d32, 9m, and p32 and for the STS parcel, we used Glasser areas STSvp, STSva, STSdp, STSda, STV. All parcels were transformed to infant-specific functional space by concatenating the subrun-to-infant template registration matrix with the infant template-to-MNI registration matrix and inverting those transformations.

We used an iterative leave-one-subrun-out procedure such that data were concatenated across all subruns except one. Then, whole-brain voxel-wise GLMs and contrast maps were computed. The top 5% of voxels that had a greater response to faces than the average response to non-face conditions within an anatomical constraint parcel were selected as the fROI for that subject. Then, the parameter estimates (i.e., beta values) for all four conditions were extracted from the left-out subrun. For all bar plots, beta values were averaged across participant sessions.

To determine whether a region’s response was category-selective, we fit the beta-values using a linear mixed effects model. In each model, we indicator-coded the three control conditions, to test the hypotheses that the response to each control condition was significantly lower than the response to the face condition. Specifically, we fit a model in R using the lme4 software package (Bates et al., 2015) with the expression:
lmer(betas ∼ bod + obj + scn + motion + age + (1|subject))

There were three indicator coded-regressors to indicate which condition the beta was from. The “bod” regressors had 1s if the beta indicated a response to bodies and 0s for betas that were from other conditions. Similarly the “obj” regressor had 1s for object betas and 0s for all other conditions and the “scn” regressor had 1s for all scene betas and 0s for all other conditions. Motion was computed as the proportion of scrubbed volumes. Age and motion were z-scored. Motion was a fixed effects parameter of no interest and subject was coded as a random effect for all models. Four individual infants contributed more than one session of data, which was treated as within-subjects variance through the random effect and the age regressor. The response in a parcel was deemed selective if the fixed effect coefficient for each of the three control conditions (i.e., bod, obj, and scn) was significantly negative. Because predictions were unidirectional, reported *p*-values were one-tailed.

We also computed (1) weighted LME models to account for the variable amount of data each subject contributed to each condition and (2) LME models with coil as a random effect. The results were similar for these additional models and are reported in Table S1.

To test for condition by age interactions, we used the following model in R:
anova(lmer(betas ∼ condName * age + motion + (1|subject)))
where condName was the condition label for each beta value; age and motion were z-scored. Post-hoc analyses of the effect of age on each condition were tested with the following model in R:
lmer(condBeta ∼ age + (1|subjID))
where condBeta was a vector with beta values from a single condition and age was z-scored. In instances where the model was singular (indicated in Table 3) we additionally fit a linear model (lm(condBeta∼zAge)) to ensure we obtained the same results.

To test for laterality effects between the left and right hemispheres, we fit the following model for each fROI in R:
anova(lmer(betas ∼ condName * hemi * age + motion + (1|subject)))
where condName was the condition label for each beta, hemisphere indicated the left or right hemisphere for each beta and motion and age were z-scored and subject was included as a random effect. We tested for effects of laterality in the approximate locations of OFA, FFA and STS, but not MPFC because the MPFC fROI is on the midline and the two hemispheres could not be reliably distinguished. For post-hoc analyses of the effect of hemisphere on each condition, we fit the following model in R for IOG, VTC, and STS:
anova(lmer(beta ∼ age * hemisphere + (1|subject)))
where beta was a vector of beta values from a single condition and hemisphere indicated whether the beta value was from the left or right hemisphere. Age was z-scored, and subject was included as a random effect.

To test if EVC has a different functional profile than face-selective regions, we fit the following model in R:
anova(lmer(betas ∼ condName * fROI + age + motion + (1|subject)))
where condName was the condition label for each beta (i.e., face, object, body, scene) and fROI was the fROI label for each beta (i.e., EVC and IOG or VTC or STS or MPFC). Motion and age were z-scored. We ran four models to test for a significant fROI by condition interaction between EVC and each face-selective fROI.

## RESULTS

### Whole Brain Contrast Maps

First, we asked if infants have face responses in the approximate locations of OFA, FFA, STS, and MPFC by visualizing whole brain maps in individual infants. At a lenient statistical threshold (*p*<0.01, uncorrected), visual inspection of contrast maps (faces>non-faces) from all sessions with sufficient usable data (n=65) revealed face activations in the approximate location of OFA, FFA, STS and MPFC in many infants scanned on both Coil 2011 and Coil 2022. Importantly, infants with different amounts of data had face responses in the approximate location of each expected region (Figure 3).

**Figure 3.**
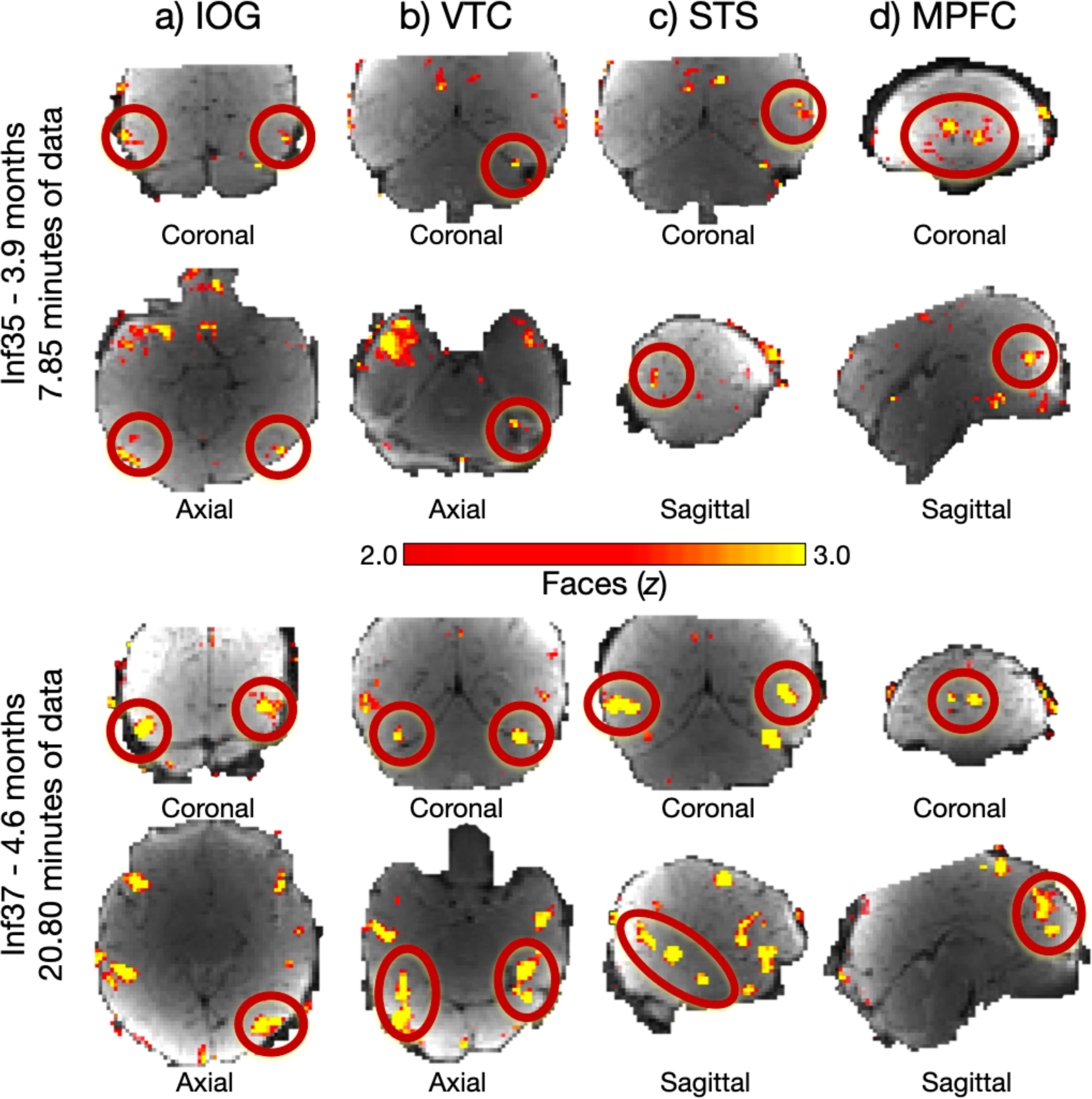
Cortical Responses to Faces in Individual Infants. Similarly aged individual infants with variable amounts of data (top panel, Inf35, 3.9 months, 7.85 minutes of data; bottom panel, Inf37, 4.6 months, 20.80 minutes of data) have activations in inferior occipital gyrus (a, IOG), the approximate location of OFA in adults, ventral temporal cortex (b, VTC), the approximate location of FFA in adults, the superior temporal cortex (c, STS), and the medial prefrontal cortex (d, MPFC). Relevant activations for each region are circled in red. Whole brain contrast maps (faces > non-faces) are displayed on a template BOLD image, truncated, and thresholded at z=2.0-3.0 to enable visualization of face activations and allow for a direct comparison of activations between the two infants. Brain orientation is indicated below each slice.

The Coil 2021 group RFX map (Figure 4) showed face>non-face activations in the approximate locations of OFA, FFA, STS, and MPFC. For Coil 2011 group RFX, there were face>non-face activations in the approximate locations of STS and MPFC but not in the approximate locations of OFA or VTC (Figure 4-1). These activations did not survive correction for multiple comparisons, though note that registration across infants was only approximate, given the highly distorted functional images, and the absence of a high resolution individual anatomical image for most individuals. Results for Coil 2011 data are visualized in Figure 4-1.

**Figure 4.**
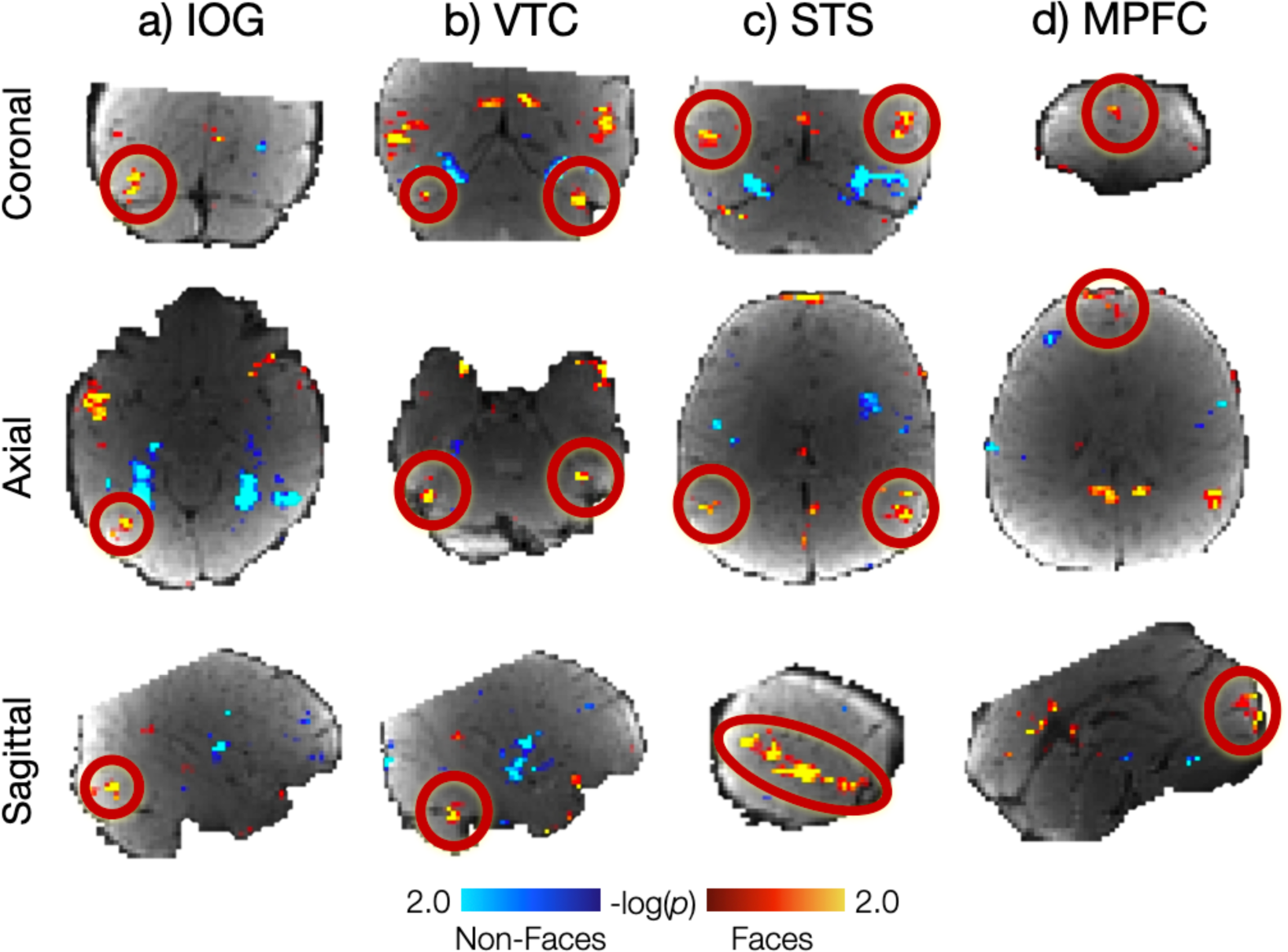
Group Face Responses in Infant Cerebral Cortex. Whole brain group random effects analysis of Coil 2021 at a lenient threshold (*p*<0.05) revealed face activations in (a) inferior occipital gyrus (IOG), the approximate location of OFA in adults, (b) ventral temporal cortex (VTC), the approximate location of FFA in adults, (c) superior temporal sulcus (STS), and (d) medial prefrontal cortex (MPFC). Hot colors indicate face activations, cool colors indicate average response to non-faces. Activations clusters did not survive correction for multiple comparisons. Activations for each region are shown on infant template BOLD image in coronal (top row), axial (middle row) and sagittal (bottom row) views and highlighted with a red circle.

### Functional Regions of Interest

All fROI analyses were conducted in n=37 sessions (33 unique individuals) who had at least two subruns. Inferior occipital gyrus (IOG) is the location of OFA in adults. Across all infants in the IOG fROI analysis (Figure 5a; Table 1) the response to faces was significantly greater than the response to bodies (*p*=0.004) and scenes (*p*<0.001) and numerically but not significantly greater than the response to objects (*p*=0.05). The overall magnitude of response across all conditions increased with age (*p*=0.006) and there was an age by condition interaction (*p*=0.02; Table 2). Post-hoc analyses revealed that the age by condition interaction was driven by a lower response to scenes in older infants (*p*=0.02; Figure 5-1a; Table 3). Using a median split by age, we ran separate fROI analyses of younger and older infants (Figure 5a; Table 1). In the younger infants alone, the response to faces was greater than the scene response (*p*=0.03) but not significantly different from the object or body responses (*ps*>0.1) but in the older infants alone the face response was significantly greater than the response to each other condition (all *ps*<0.007). To test for any laterality effects, we analyzed left and right hemispheres separately. Despite a hemisphere by age interaction (*p*=0.04; Table 4) and a trend towards a condition by age interaction (*p*=0.06), the condition by hemisphere by age interaction was not significant (*p*>0.6). Post-hoc analyses indicated there was no difference in responses to faces in right versus left hemispheres (*p*>0.4; Figure 6a; Table 4).

**Figure 5.**
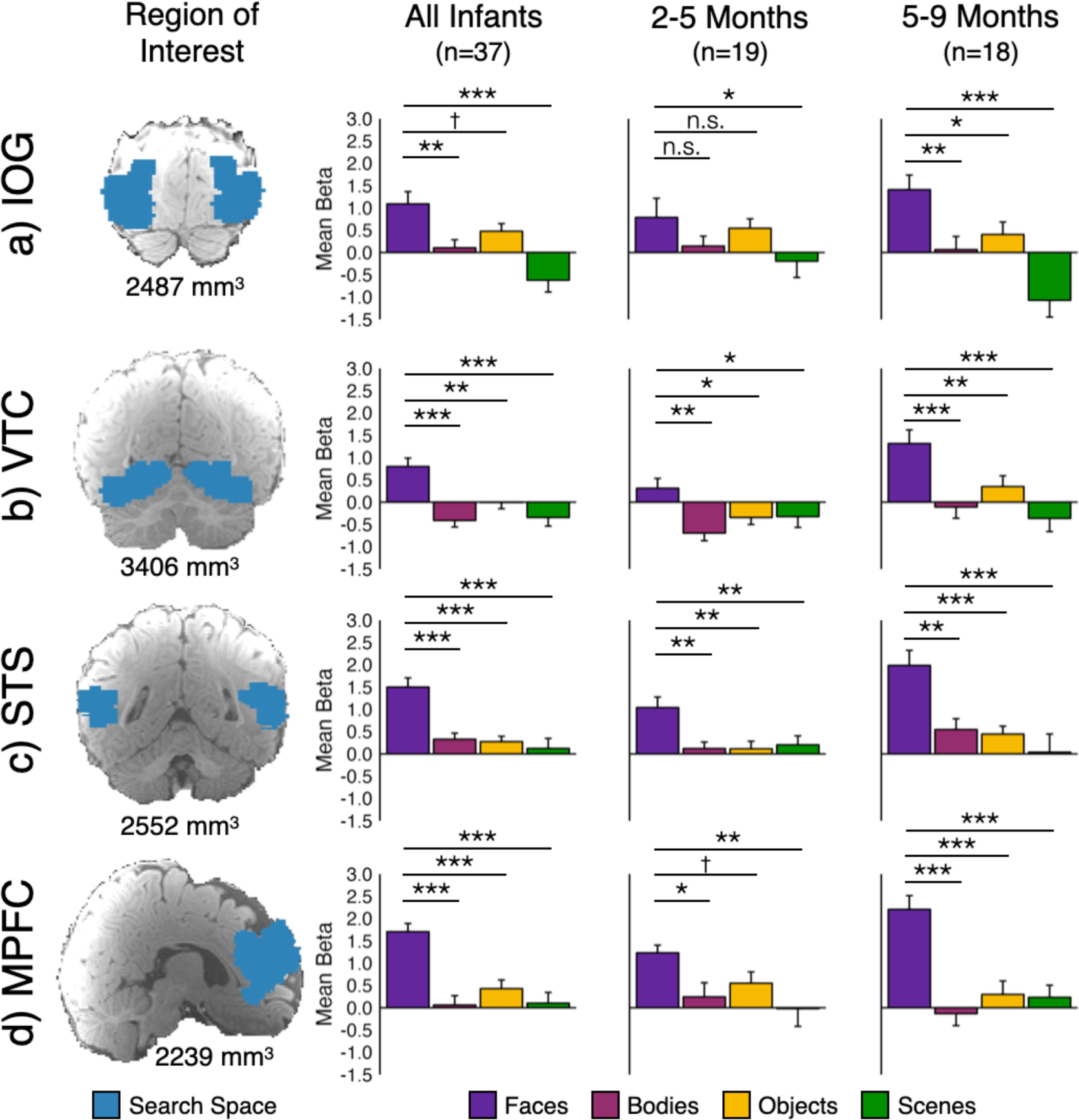
Face-Selective Responses in Infant Cortex. In analyses collapsed across Coil 2011 and Coil 2021 datasets, we identified a functional region of interest (fROI) in each participant as the top 5% of voxels that responded more to faces than non-faces within a large anatomical search space (first column, blue, projected onto an infant anatomical image) of (a) inferior occipital gyrus (IOG), the approximate location of OFA, (b) ventral temporal cortex (VTC), the approximate location of FFA, (c) superior temporal sulcus (STS), and (d) medial prefrontal cortex (MPFC). For each region, cross validated fROI analyses were conducted in all infants (n=37), and separately in younger (n=19) and older (n=18) infants. Bar charts show the mean response across participants in each fROI to faces (purple), bodies (pink), objects (yellow), and scenes (green). Error bars indicate within-subject standard error (Cousineau, 2005). Symbols indicate one-tailed statistics from linear mixed effects models: ^†^*p*<0.1; \**p*<0.05; \*\**p*<0.01; \*\*\**p*<0.001. Additional statistics reported in Tables 1 and 2; post-hoc analyses of the response to each condition as a function age are reported in Figure 5-1 and Table 3.

**Figure 6.**
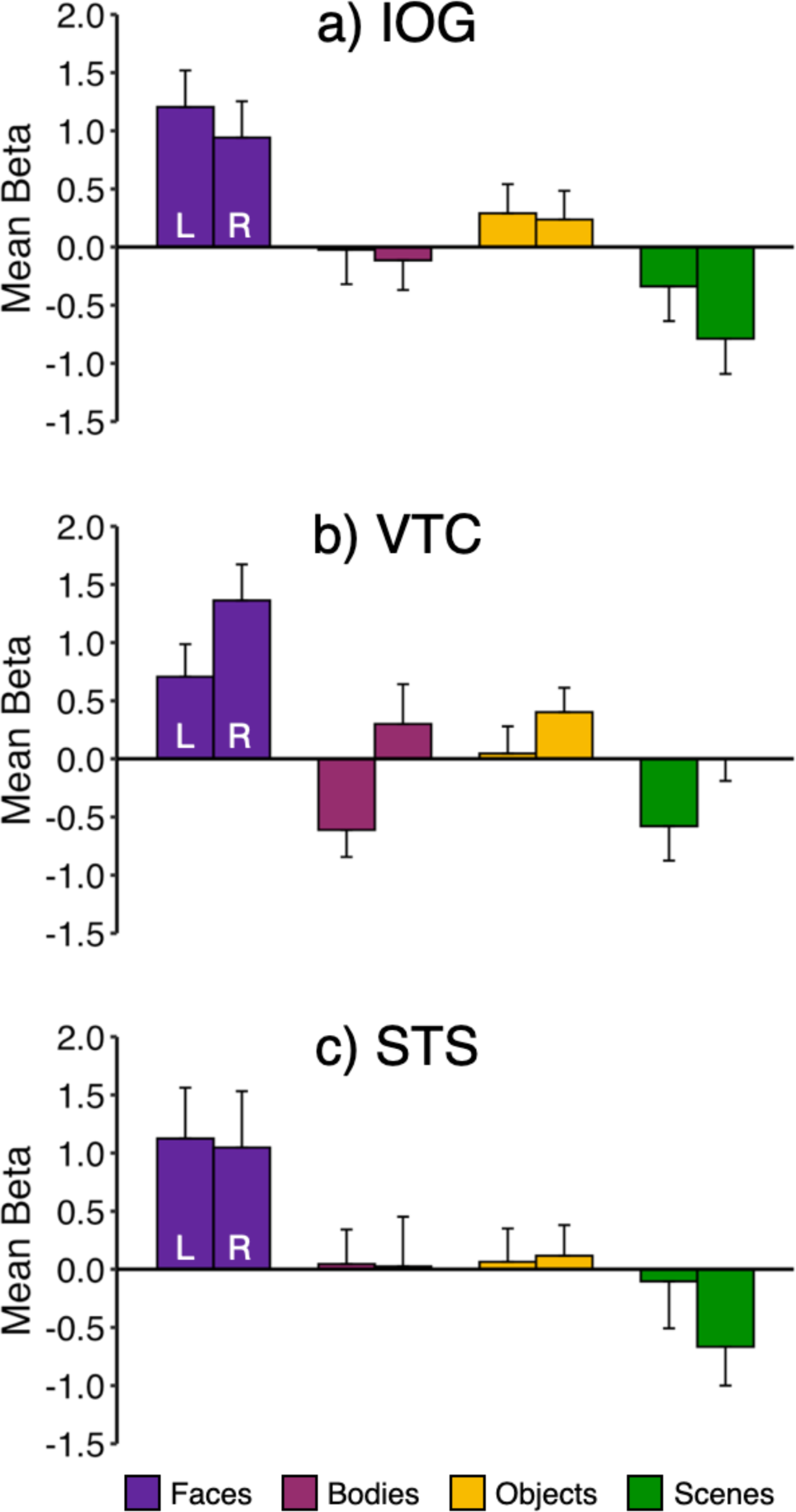
Face Selectivity is Bilateral. In analyses collapsed across Coil 2011 and Coil 2021 datasets, we identified a functional region of interest (fROI) for each participant in each hemisphere in (a) inferior occipital gyrus (IOG), the approximate location of OFA, (b) ventral temporal cortex (VTC), the approximate location of FFA, and (c) superior temporal sulcus (STS). For each region, cross validated fROI analyses were conducted in all infants (n=37). Bar charts show the mean response across participants in each fROI to faces (purple), bodies (pink), objects (yellow), and scenes (green). Error bars indicate within-subject standard error (Cousineau, 2005). Condition responses plotted as a function of age are shown in Figure 6-1; additional statistics reported in Tables 4, 4-1, 4-2, 4-3, and 4-4.

**Table 1.** Face Selectivity in Functional Regions of Interest. (Additional models in Table 1-1.)

| fROI | Intercept | Bodies | Objects | Scenes | Motion | Age |
| --- | --- | --- | --- | --- | --- | --- |
| *All Infants* |  |  |  |  |  |  |
| IOG | **1.02**<br>(0.39) | **-0.98**<br>(0.38) | *-0.61*<br>(0.38) | **-1.71**<br>(0.38) | *-0.36*<br>(0.23) | **-0.63**<br>(0.22) |
| VTC | **0.80**<br>(0.22) | **-1.21**<br>(0.29) | **-0.81**<br>(0.29) | **-1.14**<br>(0.29) | -0.15<br>(0.12) | **0.29**<br>(0.12) |
| STS | **1.51**<br>(0.26) | **-1.17**<br>(0.30) | **-1.22**<br>(0.30) | **-1.38**<br>(0.30) | 0.14<br>(0.15) | 0.13<br>(0.15) |
| MPFC | **1.66**<br>(0.33) | **-1.65**<br>(0.34) | **-1.28**<br>(0.34) | **-1.60**<br>(0.34) | -0.06<br>(0.19) | 0.12<br>(0.19) |
| EVC | -0.27<br>(0.30) | -0.32<br>(0.32) | **-0.74**<br>(0.32) | -0.16<br>(0.32) | *-0.26*<br>(0.18) | 0.09<br>(0.17) |
| *Youngest Infants* |  |  |  |  |  |  |
| IOG | **0.79**<br>(0.58) | -0.64<br>(0.52) | -0.24<br>(0.52) | **-0.98**<br>(0.52) | -0.29<br>(0.51) | 0.50<br>(0.52) |
| VTC | 0.31<br>(0.29) | **-1.01**<br>(0.33) | **-0.66**<br>(0.33) | **-0.64**<br>(0.33) | **-0.38**<br>(0.22) | 0.20<br>(0.22) |
| STS | **1.04**<br>(0.31) | **-0.92**<br>(0.31) | **-0.93**<br>(0.31) | **-0.84**<br>(0.31) | **0.21**<br>(0.25) | -0.34<br>(0.25) |
| MPFC | **1.23**<br>(0.49) | **-0.99**<br>(0.49) | *-0.68*<br>(0.49) | **-1.25**<br>(0.49) | 0.10<br>(0.42) | -0.24<br>(0.42) |
| *Oldest Infants* |  |  |  |  |  |  |
| IOG | **1.34**<br>(0.45) | **-1.24**<br>(0.52) | **-1.12**<br>(0.52) | **-2.41**<br>(0.56) | -0.25<br>(0.31) | **-0.58**<br>(0.28) |
| VTC | **1.34**<br>(0.33) | **-1.39**<br>(0.43) | **-1.17**<br>(0.43) | **-1.73**<br>(0.46) | 0.02<br>(0.19) | -0.01<br>(0.19) |
| STS | **1.98**<br>(0.39) | **-1.44**<br>(0.48) | **-1.62**<br>(0.48) | **-1.92**<br>(0.52) | 0.31<br>(0.26) | 0.14<br>(0.24) |
| MPFC | **2.13**<br>(0.41) | **-2.26**<br>(0.46) | **-2.04**<br>(0.46) | **-1.84**<br>(0.50) | 0.11<br>(0.29) | 0.32<br>(0.25) |

Parameter estimates from linear mixed effects models with beta values for each condition as predictors. Indicator-coded vectors used to test if body, object, and scene responses are each significantly less than the response to faces. Sex and z-scored age were coded as fixed effects and subject was coded as a random effect. Standard error is indicated in parentheses; $p < 0.05$ is indicated in bold; $p < 0.10$ is indicated in italics. A negative number in bold indicates a significantly lower response to that condition to faces. The intercept indicates the magnitude of the face response relative to baseline. Statistics for additional models are reported in Table 1-1.

**Table 2.** Effect of Age of Face Selectivity.

| Variable | Sum Sq. | Num. DF | Den. DF | F | P |
| --- | --- | --- | --- | --- | --- |
| **IOG** |  |  |  |  |  |
| Condition | 56.88 | 3 | 102.95 | **7.72** | **0.0001** |
| Z-Scored Age | 21.56 | 1 | 115.53 | **8.78** | **0.004** |
| Z-Scored Motion | 6.53 | 1 | 112.11 | 2.66 | 0.11 |
| Condition * Age | 24.70 | 3 | 102.95 | **3.35** | **0.02** |
| **VTC** |  |  |  |  |  |
| Condition | 34.27 | 3 | 104.00 | **7.55** | **0.0001** |
| Z-Scored Age | 8.37 | 1 | 57.12 | **5.53** | **0.02** |
| Z-Scored Motion | 2.18 | 3 | 54.11 | 1.44 | 0.23 |
| Condition * Age | 7.74 | 3 | 104.00 | 1.70 | 0.17 |
| **STS** |  |  |  |  |  |
| Condition | 44.71 | 3 | 98.72 | **9.51** | **0.00001** |
| Z-Scored Age | 1.16 | 1 | 72.50 | 0.74 | 0.39 |
| Z-Scored Motion | 0.91 | 1 | 68.93 | 0.58 | 0.45 |
| Condition * Age | 14.89 | 3 | 98.72 | **3.17** | **0.03** |
| **MPFC** |  |  |  |  |  |
| Condition | 34.27 | 3 | 104.00 | **7.55** | **0.0001** |
| Z-Scored Age | 8.37 | 1 | 57.12 | **5.53** | **0.02** |
| Z-Scored Motion | 0.18 | 1 | 97.38 | 0.08 | 0.7724 |
| Condition * Age | 7.37 | 3 | 108.35 | 1.13 | 0.3399 |

All results from linear mixed effects models converted to ANOVA table with R function anova; $p < 0.05$ is indicated in bold, $p < 0.10$ is indicated in italics.

**Table 3.** Effect of Age on Each Condition.

| **fROI** | **Intercept<sup>†</sup>** | **Age<sup>†</sup>** |
| --- | --- | --- |
| IOG Face* | **1.09 (0.48)** | 0.43 (0.48) |
| IOG Body | 0.09 (0.33) | 0.07 (0.33) |
| IOG Object | 0.45 (0.31) | -0.27 (0.31) |
| IOG Scene | *-0.64 (0.31)* | **-0.74 (0.30)** |
| VTC Face | **0.85 (0.25)** | **0.47 (0.16)** |
| VTC Body | -0.45 (0.21) | 0.32 (0.20) |
| VTC Object* | -0.01 (0.21) | 0.32 (0.21) |
| VTC Scene | -0.34 (0.22) | -0.05 (0.22) |
| STS Face* | **1.50 (0.30)** | 0.40 (0.30) |
| STS Body | 0.36 (0.26) | 0.22 (0.23) |
| STS Object | 0.30 (0.21) | 0.19 (0.19) |
| STS Scene | 0.06 (0.25) | **-0.49 (0.19)** |
| MPFC Face | **1.64 (0.35)** | 0.41 (0.31) |
| MPFC Body | -0.05 (0.37) | -0.30 (0.32) |
| MPFC Object | 0.46 (0.33) | 0.09 (0.28) |
| MPFC Scene | 0.11 (0.24) | 0.26 (0.19) |

Parameters estimated with a linear-mixed effects model in R. Condition response indicated in the left column are the predictors, z-scored age coded as a fixed effect, subject coded as a random effect. Standard error is indicated in parentheses. $p < 0.05$ is indicated in bold, $p < 0.10$ is indicated in italics. \* Indicates random effect was singular and similar results were obtained for linear model without random effect.

**Table 4.** Effect of Hemisphere on Face Selectivity.

| Variable | Sum Sq. | Num DF | Den DF | F | P |
| --- | --- | --- | --- | --- | --- |
| **IOG** |  |  |  |  |  |
| Condition | 105.06 | 3 | 240.02 | **14.58** | **0.000000009** |
| Hemisphere | 3.39 | 1 | 240.02 | 1.41 | 0.24 |
| Z-Scored Age | 0.003 | 1 | 80.05 | 0.00 | 0.97 |
| Z-Scored Motion | 7.19 | 1 | 75.17 | *2.99* | *0.09* |
| Condition * Hemi | 1.83 | 3 | 240.02 | 0.25 | 0.86 |
| Condition * Age | 20.69 | 3 | 240.02 | **2.87** | **0.04** |
| Hemisphere * Age | 8.55 | 1 | 240.02 | *3.56* | *0.06* |
| Cond * Hemi * Age | 5.66 | 3 | 240.02 | 0.78 | 0.50 |
| **VTC** |  |  |  |  |  |
| Condition | 78.71 | 3 | 251.01 | **10.67** | **0.000001** |
| Hemisphere | 28.98 | 1 | 251.01 | **11.79** | **0.0007** |
| Z-Scored Age | 40.14 | 1 | 57.07 | **16.32** | **0.0002** |
| Z-Scored Motion | 0.07 | 1 | 53.40 | 0.03 | 0.86 |
| Condition * Hemi | 2.93 | 3 | 251.01 | 0.40 | 0.75 |
| Condition * Age | 12.75 | 3 | 251.01 | 1.73 | 0.16 |
| Hemisphere * Age | 0.80 | 1 | 251.01 | 0.32 | 0.57 |
| Cond * Hemi * Age | 2.72 | 3 | 251.01 | 0.37 | 0.78 |
| **STS** |  |  |  |  |  |
| Condition | 86.44 | 3 | 246.80 | **8.06** | **0.00004** |
| Hemisphere | 1.71 | 1 | 246.80 | 0.48 | 0.49 |
| Z-Scored Age | 0.18 | 1 | 131.23 | 0.05 | 0.83 |
| Z-Scored Motion | 0.18 | 1 | 123.77 | 0.05 | 0.82 |
| Condition * Hemi | 4.33 | 3 | 246.80 | 0.40 | 0.75 |
| Condition * Age | 52.15 | 3 | 246.80 | **4.86** | **0.003** |
| Hemisphere * Age | 23.29 | 1 | 246.80 | **6.51** | **0.01** |
| Cond * Hemi * Age | 3.80 | 3 | 246.80 | 0.35 | 0.79 |

All results from linear mixed effects models converted to ANOVA with R function anova; $p < 0.05$ is indicated in bold, $p < 0.10$ is indicated in italics. Statistics for models to test for face selectivity in each hemisphere are reported in Table 4-1. ANOVA tables testing for condition by age interactions in each hemisphere are reported in Table 4-2. Statistics for models to test for effects of age on the response to each condition in each hemisphere are reported in Table 4-3. Statistics for models to test for an age by hemisphere interaction in each region are reported in Table 4-4.

Ventral temporal cortex (VTC) is the location of FFA in adults. Across all infants, in the fROI in VTC (Figure 5b; Table 1), the response to faces was significantly greater than to objects, bodies, and scenes (all *ps*<0.003). The overall magnitude of response across conditions increased with age (*p*=0.02), but there was no age by condition interaction (*p*=0.2; Table 2). However, post-hoc analyses of each condition separately (Figure 5-1; Table 3) showed that although body and object responses increase numerically with age (*ps*=0.1), only the face response was significantly greater in older infants (*p*=0.03; Figure 5-1; Table 3). In both the younger and older infants separately (Figure 5b; Table 1), the face response was significantly greater than the response to bodies (younger *p*=0.001; older *p*=0.0006), objects (younger *p*=0.02; older *p*=0.003), and scenes (younger *p*=0.03; older *p*=0.00009). The response to all stimuli was higher overall in the right hemisphere (*p*=0.0007; Table 4) and in older infants (*p*=0.0002), but there was no interaction between hemisphere and condition (*p*=0.76) or hemisphere and age (*p*=0.20).

STS contains a region that responds to both faces and voices in adults. Across all infants, in the fROI analysis of STS, the response to faces was significantly greater than to objects, bodies, and scenes (all *ps*<0.001; Figure 5c; Table 1). There was no main effect of age (*p*=0.4) but there was a condition by age interaction (*p*=0.03; Table 2) which was driven by a significantly lower response to scenes in older infants (*p*=0.03; Figure 5-1c; Table 3). In fROI analyses of the younger and older infants separately, the face response was significantly greater than responses to any other condition (all *ps*<0.03). The response to all stimuli was larger in the right hemisphere for older infants (hemisphere by age interaction, *p*=0.01) and response magnitudes were higher for older infants (age by condition interaction, *p*=0.003) but the hemisphere by age by condition interaction was not significant (*p*=0.8).

In adults, MPFC contains a region that responds to socially relevant stimuli, including faces. Across all infants, in the fROI analysis of MPFC, the response to faces was significantly greater than to objects, bodies, and scenes (all *ps*<0.001; Figure 5d; Table 1). There was no main effect of age (*p*=0.5), and no age by condition interaction (*p*=0.3; Table 2). In the younger infants alone, the MPFC face response was greater than the response to each other condition (bodies *p*=0.02, objects *p*=0.08, scenes, *p*=0.005; all *ps*<0.03 in weighted LME, Tables 1 and 1-1). Older infants also had face-selective responses in MPFC (all *ps*<0.0005) and responses to each condition did not change with age (all *ps*>0.19; Figure 5-1; Table 3).

To test whether the face-selective responses described above are spatially specific to cortical regions with face-selective responses in adults, we conducted an fROI analysis in occipital areas, the location of early visual cortex (EVC) in adults, where we do not expect to observe face-selective responses. In the fROI analysis of EVC, the face response was significantly greater than the response to objects (*p*=0.009) but was not statistically different than body and scene responses (all *ps*>0.1; Table 1). Further the face-selectivity in IOG, VTC, and MPFC fROIs, was significantly different from the absence of face selectivity in EVC (all *ps*<0.04; Table 5). The condition by fROI responses for STS versus EVC did not reach significance (*p*=0.10; Table 5). Thus, even in voxels selected for maximally face-selective responses, we confirmed that voxels in infant EVC are not face selective and are significantly different from responses in IOG, VTC, and MPFC.

**Table 5.**
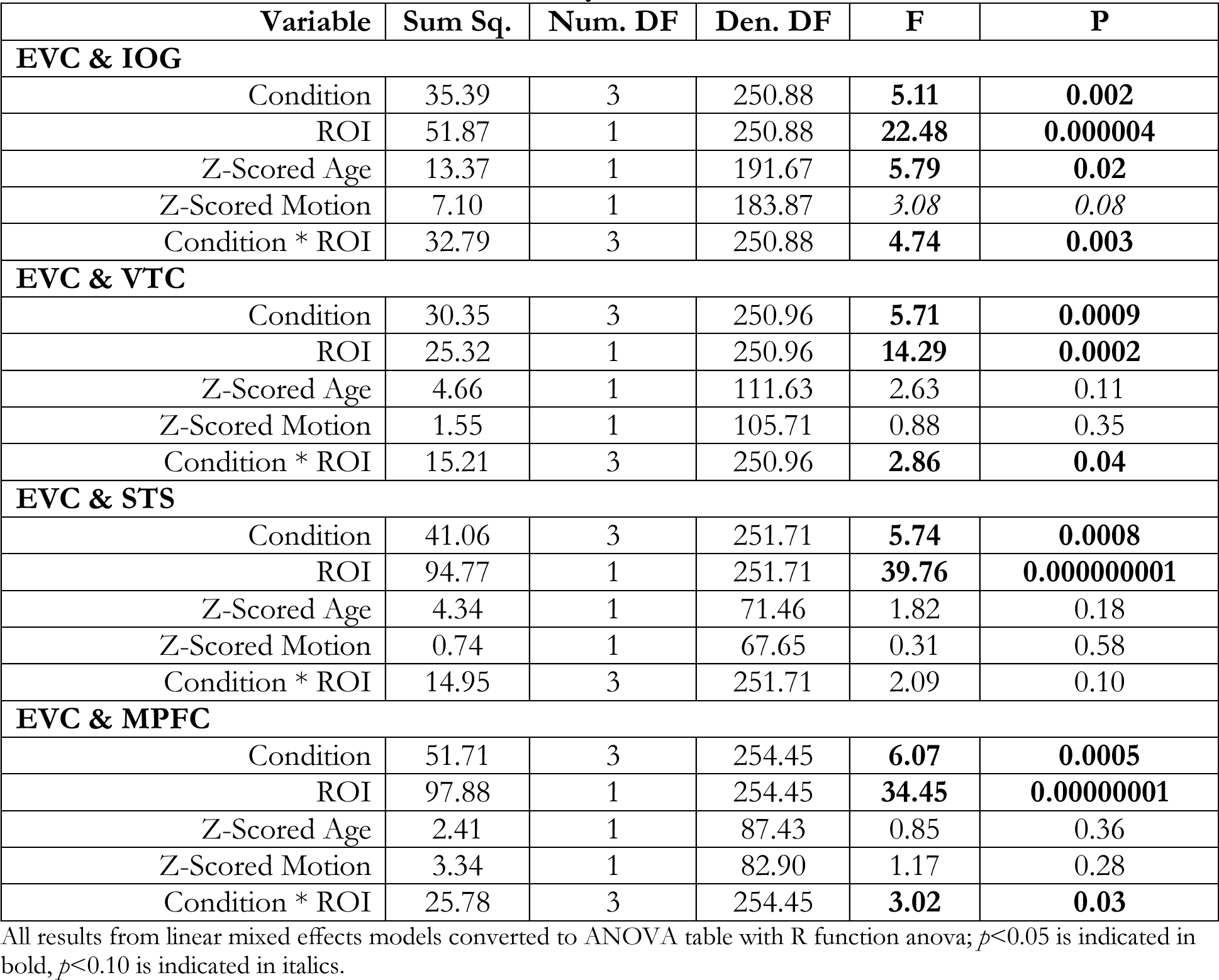
ANOVA Tables for EVC fROI analyses.

## DISCUSSION

Here we test when face-selective cortical responses first arise in the approximate location of adult OFA, FFA, STS and MPFC. We use data combined across two coils to double the sample size, compared to prior reports (Kosakowski et al., 2022a), and test whether face selectivity changes during the first year of life. To accommodate the different spatial distortions in the two coils, we used larger anatomical parcels, sacrificing spatial resolution in exchange for a larger sample of younger infants. Throughout the first year of life, including at the earliest ages we could measure, we observed face-selective responses in the approximate location of FFA, STS, and MPFC. In the approximate location of OFA, we did not observe face-selective responses in the youngest infants, but we did observe face-selective responses in older infants. Taken together, our results suggest that in humans, face-selective responses in multiple cortical regions develop in infancy.

### OFA

The region in inferior occipital gyrus, near adult OFA, appeared to show the most developmental change in our dataset. The region was face selective on average in infants and there was a significant age by condition interaction. In the older half of infants, we found face-selective responses, but not in the younger half of infants. This difference was driven a decreased response to scenes in older infants, not a change in the response to faces.

The possibility that face selectivity might arise in OFA relatively later than in FFA is intriguing, particularly because OFA was initially presumed to be the source of face-specific input for FFA (Gobbini and Haxby, 2007; Haxby et al., 2000). Yet face selectivity is still observed in FFA following focal damage to OFA (Gao et al., 2019; Rossion, 2022; Weiner et al., 2016). Thus, it is debated whether OFA is a necessary source of input to FFA in a single hierarchy, or whether FFA receives sufficient input from other sources (Pitcher, 2022; Pitcher et al., 2014; Rossion, 2022). The current results, finding face-selective responses in the youngest infants in FFA but not yet in OFA, could be construed as evidence supporting this latter idea.

However, the weaker response in the location of OFA in younger infants should be interpreted with caution. Even in adults, OFA is small, variable and difficult to detect (Dai and Scherf, 2023; Rossion et al., 2012; Schwarz et al., 2019; Zhen et al., 2015). Although we could not confirm a face-selective response near OFA in the youngest infants, we also did not find strong evidence for its absence. More data from young infants will likely be needed to establish more precisely when a face-selective response can first be detected in inferior occipital gyrus.

### FFA

Using the higher resolution subset of these data collected with Coil 2021, a previous study reported that infants have face-selective responses in the approximate location of adult FFA (Kosakowski et al., 2022a). However, the sample was too small to test for developmental change within the first year. In the current analyses of the larger sample, we find no evidence that face selectivity is late to develop. Although older infants have a greater response to faces than younger infants, the youngest infants still have a detectable face-selective response in the approximate location of FFA, and there was no age by condition interaction. The current results thus contribute to a long-standing debate about the origins of face selectivity in FFA.

In children, FFA is face selective, responding more to faces than to non-face visual categories (Aylward et al., 2005; Cantlon et al., 2011; Dehaene-Lambertz et al., 2018; Feng et al., 2022; Golarai et al., 2015, 2009, 2007; Joseph et al., 2011; Natu et al., 2016; Nordt et al., 2021; Passarotti et al., 2003; Peelen et al., 2009; Pelphrey et al., 2009; Scherf et al., 2007; Tian et al., 2021). However, compared to adults, children have (1) a smaller volume of cortex with a significant face-selective response and (2) a smaller magnitude response difference between face and non-face categories (Feng et al., 2022; Golarai et al., 2015, 2009, 2007; Haist et al., 2013; Joseph et al., 2011; Natu et al., 2016; Nordt et al., 2021; Peelen et al., 2009; Scherf et al., 2007; Tian et al., 2021) (although see (Aylward et al., 2005; Dehaene-Lambertz et al., 2018; Passarotti et al., 2003)). In children, the extent of face-selective cortex is correlated with face recognition and memory abilities (Golarai et al., 2007), which continue to develop into adulthood (Carey et al., 1992; Dundas et al., 2013; Germine et al., 2011).

Clearly, face-selective responses increase in extent and magnitude over childhood, but those observations do not establish when face-selective responses first emerge. The first neuroimaging studies with human infants suggested an early preferential response to faces in the approximate location of FFA, without full face selectivity. An early PET study demonstrated that young infants have responses to faces in the approximate location of FFA, STS, and MPFC (Tzourio-Mazoyer et al., 2002). But this study was limited because infants saw only two kinds of stimuli: static images of female faces and one control condition, colorful diodes. Similarly, an initial fMRI study found face preferences, but not selectivity, in a small sample of human infants (Deen et al., 2017). Consistent with evidence in humans, face-selective responses were observed in infant macaques only late in the first year of life (Livingstone et al., 2017), which is thought to correspond to approximately age 3 years in human development. Thus, initial PET and fMRI investigations in infants and children suggested that face selectivity in FFA might initially arise in toddlers or young children, only after substantial visual experience.

However, other fMRI and EEG data challenge that view. Two recent fMRI studies in awake infants observed face-selective responses in FFA, using different stimuli and task procedures (Kosakowski et al., 2022a; Yates et al., 2023). These results are consistent with EEG evidence of a distinctive response to faces, compared to other visual objects, in the brains of 4-to 6-month-old infants, with a source likely in VTC (De Heering and Rossion, 2015). Similarly, fMRI studies have found early origins for retinotopic organization of visual cortex (Ellis et al., 2021b; Kourtzi et al., 2006) and for responses to motion in MT (Biagi et al., 2023, 2015). Compared to these other visual regions, FFA development might be less dependent on visual experience (Li et al., 2022). Adults born with cataracts that were removed by age two, have reduced motion-related responses in MT but have preserved responses in FFA (Guerreiro et al., 2022). Further, FFA responses are heritable (Chen et al., 2023; Polk et al., 2007) and are partially preserved in congenitally blind adults (Ratan Murty et al., 2020; Van Den Hurk et al., 2017). In sum, the current results fit with a growing body of evidence that face selectivity in FFA initially arises within a few months after birth and requires little visual experience.

Still, it is likely that many aspects of the FFA response change with both age and visual experience. The initial response to faces increases with age, even in infancy and appears to expand to cover more of the fusiform gyrus during childhood (Feng et al., 2022; Golarai et al., 2015, 2009, 2007; Joseph et al., 2011; Natu et al., 2016; Nordt et al., 2021; Peelen et al., 2009; Scherf et al., 2007; Tian et al., 2021). Because of the spatial distortions we cannot confidently estimate the size of the region we observed in infants. One important question for future research will be whether the size or selectivity of FFA in infants correspond to their face recognition abilities.

### STS

A region in STS showed a robust response to faces compared to all other visual categories, in both the younger and older infants. The face stimuli were videos of children’s faces, including changing facial expressions and gaze. In adults, these videos are ideal to elicit responses in STS, which strongly prefers dynamic to static faces (Pitcher et al., 2019, 2014, 2011; Sato et al., 2004). These results converge with prior evidence of face-selective responses in the approximate location of STS in children (Pitcher et al., 2011, Walbrin et al., 2020) and, using functional near infrared spectroscopy (fNIRS), in infants (Farroni et al., 2013; Lloyd-Fox et al., 2009b; Powell et al., 2018b).

Although the STS responds selectively to faces among visual categories, in adults and older children the same region also responds to other kinds of social stimuli. For example, the face-selective region in STS responds more to point-light displays depicting two bodies interacting versus two bodies not interacting (Isik et al., 2017; Walbrin et al., 2020). The same region responds more to human speech than non-speech sounds (Deen et al., 2015). In infants, parts of STS similarly respond more to speech compared to non-speech sounds (Grossmann et al., 2010; Lloyd-Fox et al., 2014), and point-light displays depicting biological versus non-biological motion (Lisboa et al., 2020a, 2020c; Lloyd-Fox et al., 2011b). However, it is not known whether these responses are co-located in the same regions of STS as the face-selective response.

The infants studied here, as young as two months, are among the youngest in whom STS responses to dynamic faces have been reported. A limitation of the current design is that we cannot test whether this same region already also responds to human voices or other social stimuli. It is an interesting developmental question whether responses to faces and voices are processed separately in early development and then gradually associated, or whether these responses are already integrated within the first few months of life. Behaviorally, infants do seem to integrate face and voice information remarkably early. Young infants prefer to look at one face more than another based on non-visual properties (e.g., speaker language, prosody, social behavior) (Kelly et al., 2005; Kinzler et al., 2007; Turati et al., 2011). Even within the first 12 hours after birth, infants prefer to look at their own mother’s face, compared to a female stranger (Bushnell, 2001; Pascalis et al., 1995; Sai, 2005), identified by association with the mother’s voice (Sai, 2005). In one series of studies, three-hour-old infants looked more at their mother’s face, than a female stranger’s face, when they had experienced their mother’s voice and face together in those three hours; but not if their mother had been instructed to remain silent during those hours (Bushnell et al., 1989; Sai, 2005). Future research could investigate whether early developing multi-modal responses in STS are related to infants’ behavioral preferences for faces associated with specific voices.

### Laterality of Face Responses

The robust lateralization of face responses to the right hemisphere in adults (Jonas et al., 2018; Pitcher et al., 2007; Rangarajan et al., 2014; Yovel et al., 2008) was not evident in our infant data, as none of the four face-selective regions showed a significant difference between hemispheres in the profile of response across conditions. Although this result could indicate that lateralization arises later in development, it is also possible we simply lacked the power to detect effects of hemisphere (cf., the non-significant trend in infant VTC in Figure 6b). In the future, it will be important to use high-quality data from well-powered studies to determine when in development the lateralization of face selectivity in the right hemisphere emerges.

### MPFC

A region in MPFC showed face-selective responses in both younger and older infants, that did not change with age in this sample. Finding selective functional responses in MPFC in infants as young as 2-to 5-months-old is intriguing in light of the protracted anatomical development of this region (Bethlehem et al., 2022; Brody et al., 1987; Dubois et al., 2016; Kinney et al., 1988; Tau and Peterson, 2010; Vasung et al., 2019). Signatures of cortical maturation including expansion (Li et al., 2013), increased sulcal depth (Meng et al., 2014), and myelination (Carmody et al., 2004; Hasegawa et al., 1992; Miller et al., 2012) occur relatively later in MPFC than in other cortical regions. Indeed, new neurons are still migrating and being integrated into prefrontal cortex well into the second year of life (Sanai et al., 2011) – while this process is completed in primary sensory areas around the time of birth (Kostović et al., 2019).

In adults, a region in MPFC has been reported that responds more to images of faces than to other categories (Gu et al., 2023; Schwarz et al., 2019) and dynamic faces compared to dynamic objects similar to the current videos (Julian et al., 2012; Kosakowski et al., 2022b). Yet, the MPFC is not classically considered a face perception region (Gobbini and Haxby, 2007; Haxby et al., 2000) or a visual region, and its response to faces is modulated by social content (Cheng et al., 2022; LaBar, 2003; O’Doherty et al., 2003). In addition, while OFA and FFA respond only to visually presented faces, face-selective regions in MPFC also respond to a variety of other social stimuli, including animations of social interactions and stories about people presented visually and aurally (Kosakowski et al., 2022b).

Similar to adults, studies using fNIRS have reported responses to dynamic faces in infant MPFC (Farris et al., 2022; Grossmann et al., 2008a; Krol and Grossmann, 2020; Porto et al., 2020; Tzourio-Mazoyer et al., 2002). These responses are greater for socially relevant faces (e.g., a parent, direct gaze, using infant-directed speech) than faces with less social relevance (e.g., a cartoon image, an averted gaze) (Grossmann et al., 2008b; Imafuku et al., 2014; Krol and Grossmann, 2020; Lloyd-Fox et al., 2015; Naoi et al., 2012; Uchida-Ota et al., 2019; Urakawa et al., 2015; Xu et al., 2017). One intriguing possibility is that infants’ MPFC, like adult MPFC, is engaged in processing the social and emotional meaning associated with faces. However, the current evidence cannot exclude the possibility that initially infants’ MPFC contains purely visual representations of faces and only later in infancy is used to ascribe social and emotional meaning to those faces.

Our results are broadly consistent with other recent evidence of functional responses in infants’ prefrontal cortex. For example, a region in lateral prefrontal cortex in infants responds more to sequences with statistical regularity compared to unstructured input (Ellis et al., 2021a; Gervain et al., 2008; Werchan et al., 2016). Another area in infant prefrontal cortex responds more to native language compared to foreign language or other non-speech sounds (Altvater-Mackensen and Grossmann, 2018; Dehaene-Lambertz et al., 2010; May et al., 2011; Minagawa-kawai et al., 2010; Vouloumanos et al., 2010). Thus, despite structural immaturity, prefrontal cortex appears to be functionally active in infancy and may play a key role in infant cognitive development.

### Limitations and Future Directions

The current results provide an upper bound, but do not directly answer the question of when face selectivity first arises in each of the regions considered. Particularly in FFA, STS and MPFC, face-selective responses are already present in the youngest group of infants, aged 2-5 months. We cannot resolve the time of first face-selective responses more specifically than this three-month window, because we have limited data (average 16 minutes) from each infant. As a result, we cannot confidently estimate the selectivity of a region in a single infant. Moreover, we have no measurements in the first two months of infants’ lives, and so cannot determine how much earlier face-selective responses first arise. These limitations leave open the intriguing possibility that face selectivity may not arise simultaneously across these cortical regions, but instead, arise in a sequence. A strong test of this hypothesis would ideally require substantially more data per infant, collected in a dense longitudinal sample, so that the age of first face-selective responses in each region could be confidently identified.

In order to increase the number of younger infants included in this sample, we combined data across two coils. The lower resolution and more distorted images collected with Coil 2011 meant that we had to use very large parcels to identify voxels putatively near OFA and FFA in particular. Also, we did not have high resolution anatomical images for most infants. As a result, the location of the fROIs reported here is approximate. Techniques for acquiring functional data from awake infants are improving rapidly (Cusack et al., 2018; Ellis et al., 2020; Yates et al., 2021), so the current results can be replicated in the future with greater confidence in the spatial origins of the measured signals.

Finally, although each of the regions tested showed a significantly greater response to faces than the other visual categories, the actual magnitude of the face responses was very small compared to those previously measured in children and adults. There are many possible explanations of the change in hemodynamic response magnitude over development, including both changes in neural firing rates and synchrony (Kiorpes, 2015; Uhlhaas et al., 2010), and changes in vasculature and neurovascular coupling (Colonnese et al., 2008). The current study cannot differentiate between these explanations. However, the change in magnitude we observed between age 2 and 9 months was gradual and moderate, consistent with other evidence that the magnitude of hemodynamic responses change slowly and gradually throughout infancy and childhood (Arichi et al., 2012; Cohen Kadosh et al., 2013, 2011; Cusack et al., 2015; De Oliveira et al., 2017).

### Summary

In sum, using fMRI data from a large sample of awake infants, we measured face-selective responses in multiple regions of infants’ brains. Robust face selectivity was present in the approximate location of FFA, STS, and MPFC as early as we could measure. Putative OFA also had face-selective responses in older infants but not yet younger infants. Despite undergoing rapid anatomical change in the first postnatal year, infant cortex already has structured responses to meaningful, self-relevant stimuli such as faces. These results importantly constrain theories of cortical development and the origins of face selectivity.

## Acknowledgments

This research was carried out at the Athinoula A. Martinos Imaging Center at the McGovern Institute for Brain Research at MIT. The authors thank Nayanika Das and Somaia Saba for help with registrations; Steven Shannon, Atsushi Takahashi, and Boris Keil for technical support; members of Saxe Lab, and members of the Kanwisher Lab for help during recruitment and data collection; the Cambridge Writing Group, and members of the Saxe Lab and Kanwisher Lab for helpful comments on various version of the manuscript; Michelle Hung and Kirsten Lydic for code review; Sofia Riskin for data reconciliation; Hannah LeBlanc for all the things; and all the infants and their families. **Funding:** We gratefully acknowledge support of this project by a National Science Foundation (graduate fellowship to HLK; Collaborative Research Award #1829470 to MAC), NIH (#1F99NS124175 to HLK; #8K00DA058542-02 to HLK; #R21-HD090346-02 to RS; #DP1HD091947 to NK; shared instrumentation grant S10OD021569 for the MRI scanner), Templeton World Charity Foundation (#2022-30268 & #2022-30269 to MAC), Canadian Institute for Advanced Research Azrieli Global Scholars Fellowship (to MAC), the McGovern Institute for Brain Research at MIT, and the Center for Brains, Minds and Machines (CBMM), funded by an NSF STC award (CCF-1231216).

## Author contributions

HLK and RS designed the study. HLK, LH, IN, and MAC collected the data. HLK analyzed the data with input from and supervision by MAC, NK, and RS. HLK, NK, and RS wrote the manuscript. All authors provided feedback on the final version.

