## Supplemental Tables and Figures for "Cortical Face-Selective Responses Emerge Early in Human Infancy"

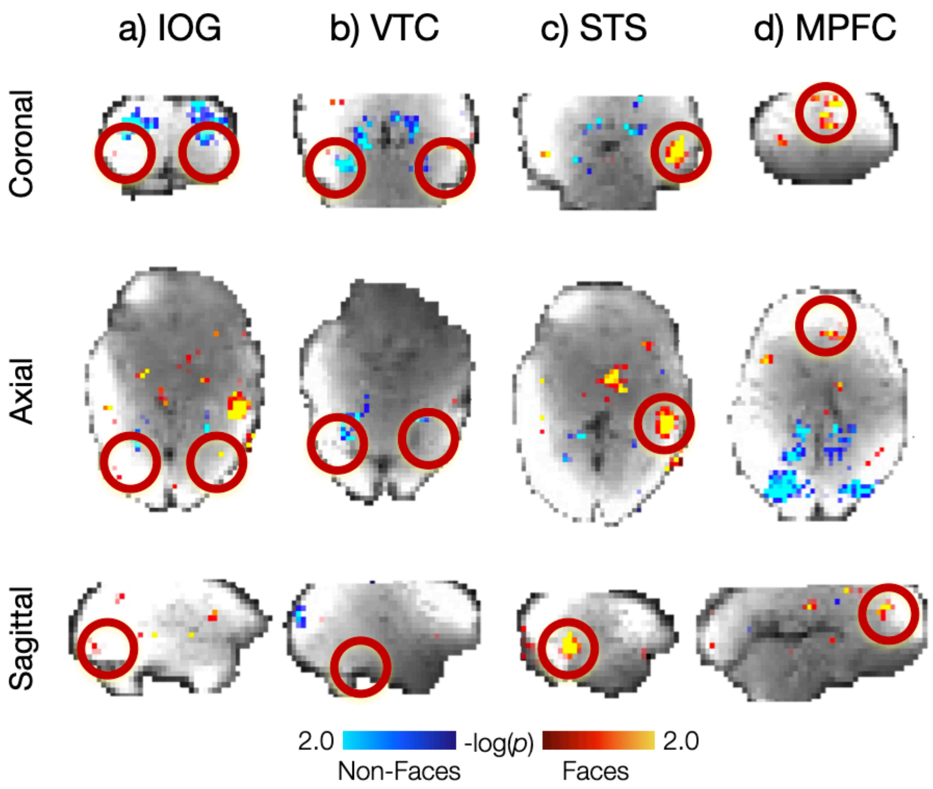


**Figure 4-1. Group Face Responses in Infant Cerebral Cortex.** Whole brain group random effects analysis of Coil 2011 at a lenient threshold ($p<0.05$) revealed face activations in (c) superior temporal sulcus (STS), and (d) medial prefrontal cortex (MPFC). Face activations were not observed in the group in (a) inferior occipital gyrus (IOG), the approximate location of OFA in adults or (b) ventral temporal cortex (VTC), the approximate location of FFA in adults. Hot colors indicate face activations, cool colors indicate average response to non-faces. Activation clusters did not survive correction for multiple comparisons. Activations for each region are shown on infant template BOLD image in coronal (top row), axial (middle row) and sagittal (bottom row) views and highlighted with a red circle. Results for Coil 2021 data are visualized in Figure 4.

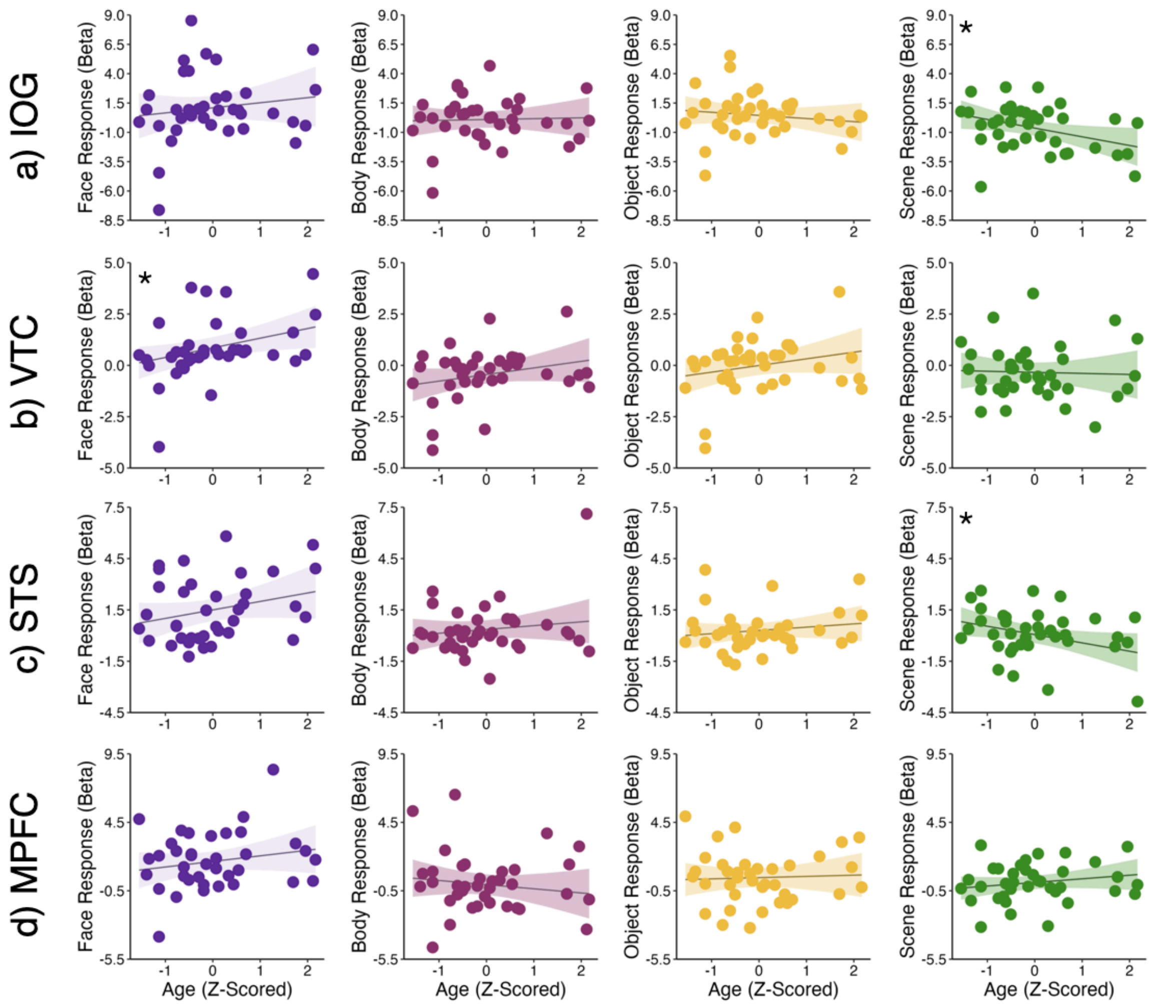


**Figure 5-1. Effect of Age on the Response Magnitude for Each Condition in Each fROI.** Scatter polots show magnitudes for each condition from fROI analyses collapsed across Coil 2011 and Coil 2021 datasets as a function of age. ROIs include (a) inferior occipital gyrus (IOG), the approximate location of OFA, (b) ventral temporal cortex (VTC), the approximate location of FFA, (c) superior temporal sulcus (STS), and (d) MPFC. Face betas are plotted in purple, body betas are plotted in pink, object betas are plots in yellow, and scene betas are plotted in green. Age is z-scored. Symbols indicate statistics from linear mixed effects models: $^{*}p<0.05$. Additional statistics reported in Table 3.

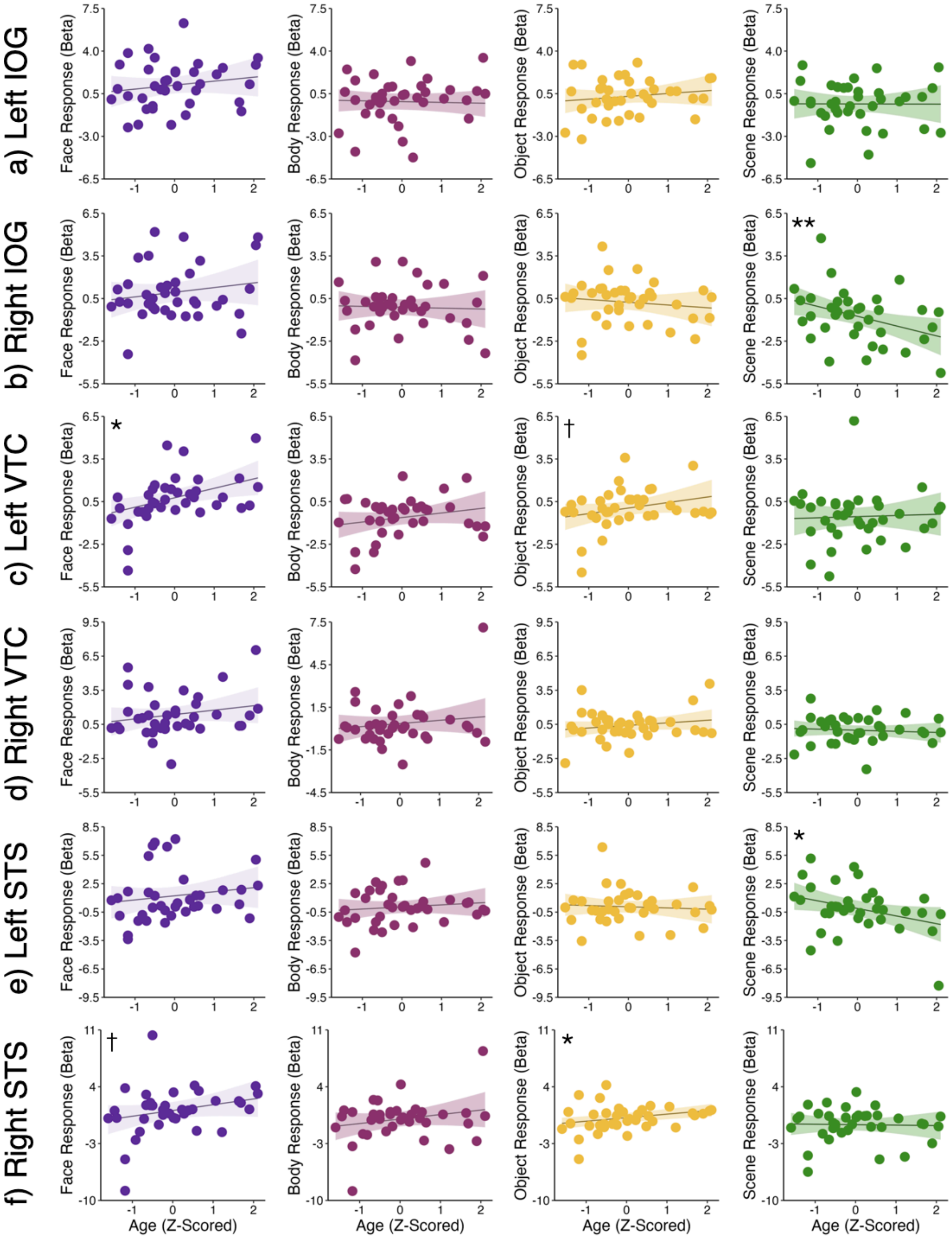


**Figure 6-1. Effect of Age on the Response Magnitude for Each Condition in Each fROI.** Scatter plots show magnitudes for each condition from fROI analyses collapsed across Coil 2011 and Coil 2021 datasets as a function of age. ROIs include (a, b) left and right inferior occipital gyrus (IOG), the approximate location of OFA, (c, d) left and right ventral temporal cortex (VTC), the approximate location of FFA, and (e, f) superior temporal sulcus (STS). Age is z-scored. Face betas are plotted in purple, body betas are plotted in pink, object

betas are plots in yellow, and scene betas are plotted in green. Symbols indicate statistics from linear mixed effects models: †$p<0.1$; *$p<0.05$; **$p<0.01$. Additional statistics reported in Table 4-2.

**Table 1-1. Additional Models for Face-Selective Responses in fROIs.**

| **fROI Analysis** | **Intercept†** | **Bodies†** | **Objects†** | **Scenes†** | **Motion†** | **Age†** |
|---|---|---|---|---|---|---|
| ***All Infants*** | | | | | | |
| IOG (w) | **1.30 (0.38)** | **-1.35 (0.39)** | **-0.83 (0.36)** | **-1.98 (0.38)** | -0.37 (0.23) | **-0.67 (0.22)** |
| VTC (w) | **0.90 (0.20)** | **-1.28 (0.28)** | **-0.78 (0.26)** | **-1.25 (0.27)** | -0.08 (0.13) | **0.31 (0.12)** |
| STS (w) | **1.52 (0.24)** | **-1.15 (0.28)** | **-1.20 (0.26)** | **-1.36 (0.27)** | -0.02 (0.15) | 0.09 (0.15) |
| MPFC (w) | **1.65 (0.30)** | **-1.59 (0.32)** | **-1.26 (0.30)** | **-1.44 (0.31)** | -0.14 (0.19) | 0.17 (0.18) |
| EVC (w) | -0.14 (0.26) | *-0.45 (0.31)* | **-0.97 (0.28)** | -0.26 (0.30) | -0.23 (0.17) | 0.12 (0.16) |
| IOG (CR) | **1.02 (0.39)** | **-0.98 (0.38)** | *-0.61 (0.38)* | **-1.71 (0.38)** | *-0.36 (0.23)* | **-0.63 (0.22)** |
| VTC (CR) | **0.81 (0.24)** | **-1.21 (0.29)** | **-0.81 (0.29)** | **-1.14 (0.29)** | -0.15 (0.12) | **0.30 (0.12)** |
| STS (CR) | **1.51 (0.28)** | **-1.17 (0.30)** | **-1.22 (0.30)** | **-1.38 (0.30)** | 0.14 (0.15) | 0.12 (0.15) |
| MPFC (CR) | **1.66 (0.33)** | **-1.65 (0.34)** | **-1.28 (0.34)** | **-1.60 (0.34)** | **-0.06 (0.19)** | 0.12 (0.19) |
| EVC (CR) | -0.22 (0.34) | -0.32 (0.31) | -0.74 (0.31) | -0.16 (0.31) | -0.27 (0.18) | 0.12 (0.18) |
| ***Youngest Infants*** | | | | | | |
| IOG (w) | **1.24 (0.59)** | **-1.17 (0.58)** | -0.61 (0.55) | **-1.40 (0.56)** | -0.27 (0.53) | 0.55 (0.53) |
| VTC (w) | *0.45 (0.27)* | **-1.15 (0.33)** | **-0.63 (0.31)** | **-0.85 (0.32)** | *-0.37 (0.22)* | 0.21 (0.22) |
| STS (w) | **1.14 (0.30)** | **-1.04 (0.32)** | **-1.02 (0.30)** | **-0.94 (0.30)** | 0.21 (0.27) | -0.37 (0.27) |
| MPFC (w) | **1.28 (0.44)** | **-1.15 (0.40)** | **-0.70 (0.38)** | **-1.05 (0.39)** | 0.05 (0.41) | -0.19 (0.41) |
| IOG (CR) | *0.79 (0.59)* | -0.64 (0.52) | -0.24 (0.52) | **-0.98 (0.52)** | -0.28 (0.52) | 0.50 (0.52) |
| VTC (CR) | 0.34 (0.33) | **-1.01 (0.33)** | **-0.66 (0.33)** | **-0.64 (0.33)** | *-0.39 (0.22)* | 0.20 (0.21) |
| STS (CR) | 1.04 (0.31) | -0.92 (0.31) | -0.93 (0.31) | -0.84 (0.31) | 0.21 (0.25) | -0.34 (0.25) |
| MPFC (CR) | **1.27 (0.55)** | **-1.00 (0.49)** | *-0.68 (0.49)* | **-1.25 (0.49)** | 0.01 (0.43) | -0.24 (0.41) |
| ***Oldest Infants*** | | | | | | |
| IOG (w) | **1.46 (0.42)** | **-1.44 (0.50)** | **-1.13 (0.46)** | **-2.72 (0.55)** | -0.19 (0.33) | *-0.54 (0.27)* |
| VTC (w) | **1.44 (0.30)** | **-1.51 (0.38)** | **-1.18 (0.35)** | **-1.93 (0.41)** | 0.05 (0.23) | 0.01 (0.19) |
| STS (w) | **1.97 (0.34)** | **-1.37 (0.41)** | **-1.59 (0.38)** | **-1.84 (0.45)** | 0.29 (0.26) | 0.20 (0.22) |
| MPFC (w) | **2.05 (0.42)** | **-1.95 (0.49)** | **-1.87 (0.44)** | **-1.73 (0.52)** | 0.14 (0.34) | 0.36 (0.27) |
| IOG (CR) | **1.34 (0.50)** | **-1.23 (0.53)** | **-1.11 (0.53)** | **-2.43 (0.57)** | -0.25 (0.30) | **-0.58 (0.29)** |
| VTC (CR) | **1.34 (0.33)** | **-1.39 (0.43)** | **-1.17 (0.43)** | **-1.73 (0.46)** | 0.02 (0.19) | -0.01 (0.19) |
| STS (CR) | **1.98 (0.39)** | **-1.44 (0.48)** | **-1.62 (0.48)** | **-1.92 (0.52)** | 0.31 (0.26) | 0.14 (0.24) |

| | | | | | | |
|---|---|---|---|---|---|---|
| MPFC (CR) | **2.13 (0.41)** | **-2.26 (0.46)** | **-2.04 (0.46)** | **-1.84 (0.50)** | 0.11 (0.29) | 0.32 (0.25) |

Parameter estimates from linear mixed effects models with beta values for each condition as predictors, (w) indicates a weight vector was included in the model, (CR) indicates coil was included as a random effects regressor in the model (see methods). Indicator-coded vectors used to test if body, object, and scene responses are each significantly less than the response to faces. Sex and z-scored age were coded as fixed effects and subject was coded as a random effect. Standard error is indicated in paratheses; $p<0.05$ is indicated in bold; $p<0.10$ is indicated in italics.

**Table 4-1. Effect of Age on Each Condition in Each Hemisphere.**

| **fROI** | **Intercept** | **Bodies** | **Objects** | **Scenes** | **Motion** | **Age** |
|---|---|---|---|---|---|---|
| Left IOG | **1.14 (0.31)** | **-1.23 (0.31)** | **-0.91 (0.31)** | **-1.54 (0.31)** | -0.25 (0.18) | -0.06 (0.18) |
| Right IOG | **0.92 (0.28)** | **-1.06 (0.36)** | **-0.70 (0.36)** | **-1.73 (0.36)** | *-0.29 (0.16)* | -0.26 (0.16) |
| Left VTC | **0.72 (0.26)** | **-1.32 (0.32)** | **-0.66 (0.32)** | **-1.29 (0.32)** | *-0.25 (0.15)* | **0.36 (0.15)** |
| Right VTC | **1.36 (0.28)** | **-1.06 (0.28)** | **-0.96 (0.28)** | **-1.36 (0.28)** | 0.12 (0.16) | 0.19 (0.16) |
| Left STS | **1.17 (0.38)** | **-1.08 (0.43)** | **-1.06 (0.43)** | **-1.23 (0.43)** | -0.22 (0.22) | -0.04 (0.22) |
| Right STS | **1.09 (0.42)** | **-1.02 (0.39)** | **-0.93 (0.39)** | **-1.72 (0.39)** | 0.25 (0.24) | 0.10 (0.24) |

Parameter estimates from linear mixed effects models with beta values for each condition as predictors. Indicator-coded vectors used to test if body, object, and scene responses are each significantly less than the response to faces. Sex and z-scored age were coded as fixed effects and subject was coded as a random effect. Standard error is indicated in paratheses; $p<0.05$ is indicated in bold; $p<0.10$ is indicated in italics.

**Table 4-2. Effect of Age of Face Selectivity in Each Hemisphere.**

| Variable | Sum Sq. | Num. DF | Den. DF | F | P |
|---|---|---|---|---|---|
| **Left IOG** | | | | | |
| Condition | 49.24 | 3 | 104.56 | **9.10** | **0.00002** |
| Z-Scored Age | 0.21 | 1 | 102.49 | 0.11 | 0.74 |
| Z-Scored Motion | 3.45 | 1 | 100.14 | 1.91 | 0.17 |
| Condition * Age | 2.90 | 3 | 104.56 | 0.54 | 0.66 |
| **Right IOG** | | | | | |
| Condition | 57.64 | 3 | 106.23 | **8.60** | **0.00004** |
| Z-Scored Age | 5.97 | 1 | 67.02 | 2.67 | 0.11 |
| Z-Scored Motion | 7.41 | 1 | 64.47 | *3.31* | *0.07* |
| Condition * Age | 18.76 | 3 | 106.23 | **2.80** | **0.04** |
| **Left VTC** | | | | | |
| Condition | 43.06 | 3 | 106.97 | **7.64** | **0.0001** |
| Z-Scored Age | 11.66 | 1 | 67.33 | **6.21** | **0.02** |
| Z-Scored Motion | 5.40 | 1 | 64.76 | *2.88* | *0.09* |
| Condition * Age | 9.82 | 3 | 106.97 | 1.74 | 0.16 |
| **Right VTC** | | | | | |
| Condition | 38.59 | 3 | 104.72 | **8.95** | **0.00002** |
| Z-Scored Age | 2.02 | 1 | 105.66 | 1.40 | 0.24 |
| Z-Scored Motion | 0.72 | 1 | 103.37 | 0.50 | 0.48 |
| Condition * Age | 4.80 | 3 | 104.72 | 1.11 | 0.35 |
| **Left STS** | | | | | |
| Condition | 35.66 | 3 | 106.68 | **3.70** | **0.01** |
| Z-Scored Age | 0.11 | 1 | 86.49 | 0.03 | 0.85 |
| Z-Scored Motion | 3.54 | 1 | 84.05 | 1.10 | 0.30 |
| Condition * Age | 38.21 | 3 | 106.68 | **3.97** | **0.01** |
| **Right STS** | | | | | |
| Condition | 55.11 | 3 | 102.85 | **6.63** | **0.0004** |
| Z-Scored Age | 0.45 | 1 | 114.68 | 0.16 | 0.69 |
| Z-Scored Motion | 3.17 | 1 | 112.58 | 1.15 | 0.29 |
| Condition * Age | 15.96 | 3 | 102.85 | 1.92 | 0.13 |

All results from linear mixed effects models converted to ANOVA table with R function anova; $p<0.05$ is indicated in bold, $p<0.10$ is indicated in italics.

**Table 4-3 Effects of Age on Each Condition in Each Hemisphere.**

| fROI | Left Intercept | Left Age | Right Intercept | Right Age |
|---|---|---|---|---|
| IOG Face | **1.20 (0.31)** | 0.33 (0.31) | **0.94 (0.31)** | 0.33 (0.31) |
| IOG Body | -0.15 (0.32) | -0.06 (0.17) | -0.11 (0.26) | -0.06 (0.26) |
| IOG Object | 0.28 (0.25) | 0.23 (0.25) | 0.24 (0.25) | -0.19 (0.25) |
| IOG Scene | -0.34 (0.30) | -0.01 (0.30) | **-0.75 (0.29)** | **-0.69 (0.22)** |
| VTC Face | **0.77 (0.27)** | **0.66 (0.20)** | **1.36 (0.31)** | 0.39 (0.31) |
| VTC Body | **-0.62 (0.23)** | 0.32 (0.23) | 0.30 (0.34) | 0.49 (0.34) |
| VTC Object | 0.05 (0.23) | *0.39 (0.23)* | *0.41 (0.22)* | 0.23 (0.15) |
| VTC Scene | *-0.56 (0.31)* | 0.08 (0.28) | -0.02 (0.21) | -0.10 (0.15) |
| STS Face | **1.22 (0.46)** | 0.43 (0.39) | **1.08 (0.50)** | *0.71 (0.38)* |
| STS Body | 0.05 (0.30) | 0.22 (0.30) | 0.03 (0.42) | 0.55 (0.42) |
| STS Object | 0.06 (0.29) | -0.13 (0.29) | 0.19 (0.28) | **0.43 (0.10)** |
| STS Scene | -0.12 (0.41) | **-0.80 (0.31)** | *-0.67 (0.34)* | -0.05 (0.34) |

Parameters estimated with a linear-mixed effects model in R. Condition responses are the predictors, age is z-scored and coded as a fixed effect and subject was coded as a random effect. Standard error is indicated in paratheses. $p$<0.05 is indicated in bold, $p$<0.10 is indicated in italics. * Indicates random effect was singular but similar results were obtained for linear model without random effect.

**Table 4-3. Effect of Hemisphere on Each Condition.**

| Variable | Sum Sq. | Num DF | Den DF | F | P |
|---|---|---|---|---|---|
| **IOG - Face** | | | | | |
| Z-Scored Age | 2.34009 | 1 | 61.071 | 0.9341 | 0.33 |
| Hemisphere | 1.28017 | 1 | 35.089 | 0.5110 | 0.48 |
| Hemisphere * Age | 0.00008 | 1 | 35.089 | 0.00 | 0.96 |
| **IOG - Body** | | | | | |
| Z-Scored Age | 0.86999 | 1 | 53.461 | 0.3401 | 0.5622 |
| Hemisphere | 0.15073 | 1 | 37.976 | 0.0589 | 0.8095 |
| Hemisphere * Age | 3.01633 | 1 | 37.976 | 1.1791 | 0.2844 |
| **IOG - Object** | | | | | |
| Z-Scored Age | 0.2401 | 1 | 58.801 | 0.1389 | 0.7107 |
| Hemisphere | 0.0528 | 1 | 36.639 | 0.0305 | 0.8622 |
| Hemisphere * Age | 3.4915 | 1 | 36.639 | 2.0202 | 0.1637 |
| **IOG - Scene** | | | | | |
| Z-Scored Age | 6.9985 | 1 | 58.924 | 2.7845 | 0.10049 |
| Hemisphere | 3.7284 | 1 | 39.913 | 1.4834 | 0.23039 |
| Hemisphere * Age | 8.0793 | 1 | 39.913 | *3.2145* | *0.08057* |
| **VTC - Face** | | | | | |
| Z-Scored Age | 24.9772 | 1 | 70 | **8.5872** | **0.004568** |
| Hemisphere | 7.9593 | 1 | 70 | 2.7364 | 0.102562 |
| Hemisphere * Age | 2.7933 | 1 | 70 | 0.9603 | 0.330477 |
| **VTC - Body** | | | | | |
| Z-Scored Age | 12.003 | 1 | 70 | *3.9177* | *0.05172* |
| Hemisphere | 15.410 | 1 | 70 | **5.0297** | **0.02808** |
| Hemisphere * Age | 0.561 | 1 | 70 | 0.1831 | 0.67002 |
| **VTC - Object** | | | | | |
| Z-Scored Age | 7.5515 | 1 | 70 | **4.2594** | **0.04274** |
| Hemisphere | 2.3334 | 1 | 70 | 1.3162 | 0.25519 |
| Hemisphere * Age | 0.3614 | 1 | 70 | 0.2039 | 0.65301 |
| **VTC - Scene** | | | | | |
| Z-Scored Age | 0.0760 | 1 | 49.747 | 0.0335 | 0.8556 |
| Hemisphere | 6.2084 | 1 | 36.675 | 2.7344 | 0.1067 |
| Hemisphere * Age | 0.0412 | 1 | 36.675 | 0.0182 | 0.8935 |
| **STS - Face** | | | | | |
| Z-Scored Age | 7.6681 | 1 | 68.812 | *2.9554* | *0.09009* |
| Hemisphere | 0.1110 | 1 | 37.760 | 0.0428 | 0.83727 |
| Hemisphere * Age | 3.5132 | 1 | 37.760 | 1.3540 | 0.25188 |
| **STS - Body** | | | | | |
| Z-Scored Age | 7.7393 | 1 | 55.084 | 1.8512 | 0.1792 |
| Hemisphere | 0.0071 | 1 | 37.023 | 0.0017 | 0.9673 |
| Hemisphere * Age | 2.0582 | 1 | 37.023 | 0.4923 | 0.4873 |
| **STS - Object** | | | | | |
| Z-Scored Age | 1.4163 | 1 | 61.989 | 0.7383 | 0.3935 |
| Hemisphere | 0.0509 | 1 | 36.875 | 0.0265 | 0.8716 |
| Hemisphere * Age | 5.0292 | 1 | 36.875 | 2.6216 | 0.1139 |

| **STS - Scene** | | | | | |
|---|---|---|---|---|---|
| Z-Scored Age | 15.1714 | 1 | 55.573 | *3.9801* | *0.05095* |
| Hemisphere | 5.8709 | 1 | 34.798 | 1.5402 | 0.22289 |
| Hemisphere * Age | 13.6255 | 1 | 34.798 | *3.5746* | *0.06702* |

Parameters estimated with a linear-mixed effects model in R. Condition response indicated in the left column are the predictors, z-scored age coded as a fixed effect, hemisphere coded as a fixed effect, subject coded as a random effect. Standard error is indicated in paratheses. $p<0.05$ is indicated in bold, $p<0.10$ is indicated in italics.
